# Cannabidiol (CBD) Promotes Post-TBI Astrocyte Viability and Decreases Injury-Induced Glial Stress Responses Across Zebra Finch Song Control Nuclei

**DOI:** 10.64898/2026.04.13.717820

**Authors:** Dylan A. Marshall, Karen A. Litwa, Ken Soderstrom

## Abstract

The non-euphorigenic phytocannabinoid cannabidiol (CBD) has demonstrated therapeutic efficacy in childhood-onset epilepsies. Using a songbird preclinical model we have found that CBD promotes recovery of learned vocalizations following focal motor cortical injury. But questions about cellular mechanisms supporting this protection remained. Songbird vocal learning, like human speech, depends on development and maintenance of specialized neural circuits. Partial lesioning (microlesions) of the vocal pre-motor cortical-like song region HVC transiently disrupts song structure and triggers injury-associated cellular stress responses across interconnected song regions. Building on prior findings that CBD reduces neuroinflammation and synaptic loss in zebra finch song circuitry, we investigated potential astrocyte contributions. Here we report that HVC microlesions induce significant cell loss in HVC and its projection targets (vocal motor RA and striatal Area X), with a substantial fraction of apoptotic cells being astrocytes. CBD treatment reduces lesion-induced apoptosis and preserves astrocyte populations, indicating enhanced astrocyte viability as a major factor in CBD-mediated neuroprotection. Microlesions also elevate astrocyte stress, including increased lysosomal burden (LAMP1/LC3 expression) and astrocytic reactivity markers (C3, S100A10, aromatase). CBD attenuates these stress responses while enhancing neuroprotective metabolic and antioxidant mediators (glutamine synthetase [GS], glutamate-cysteine ligase modifier subunit [GCLM]), consistent with improved antioxidant and excitotoxicity resistance. Given that development-dependent sensorimotor skills (e.g. song in songbirds, language and many others in humans) depend on sensitive period establishment and ongoing post-learning maintenance of specialized neural circuits vulnerable to traumatic disruption, the zebra finch model provides a valuable preclinical platform for investigating glial-targeted interventions to promote circuit resilience and functional recovery after TBI.

## Introduction

Cannabidiol (CBD), a non-euphorigenic phytocannabinoid, effectively treats childhood-onset epilepsies often linked to speech/language delays [1–5]. This motivates exploration of CBD’s influence on neural circuits underlying vocal/sensorimotor communication. Unlike Δ⁹-tetrahydrocannabinol (THC), CBD exhibits broad neuroprotective activity through diverse targets [6–8] positioning it as a candidate for studying circuit protection and/or recovery following CNS trauma.

Vocal learning, a conserved trait in humans and select species [9] is modeled in songbirds like the zebra finch. Song is acquired during a sensitive developmental period and maintained/refined via lifelong sensorimotor feedback [10]. The song system includes discrete cortical-like (HVC), motor (RA), and basal ganglia/striatal (Area X) nuclei [11]. HVC, a vocal premotor circuit node, sequences syllables/syntax and integrates auditory input [12–14]. HVC projects to RA for vocal motor output and Area X (basal ganglia homolog) for variability/plasticity via the anterior forebrain pathway (AFP) to RA (Mooney, 2009, See Fig. 1). Area X uniquely integrates striatal and pallidal features in one nucleus, enabling input/output modulation [15].

**Figure 1:**
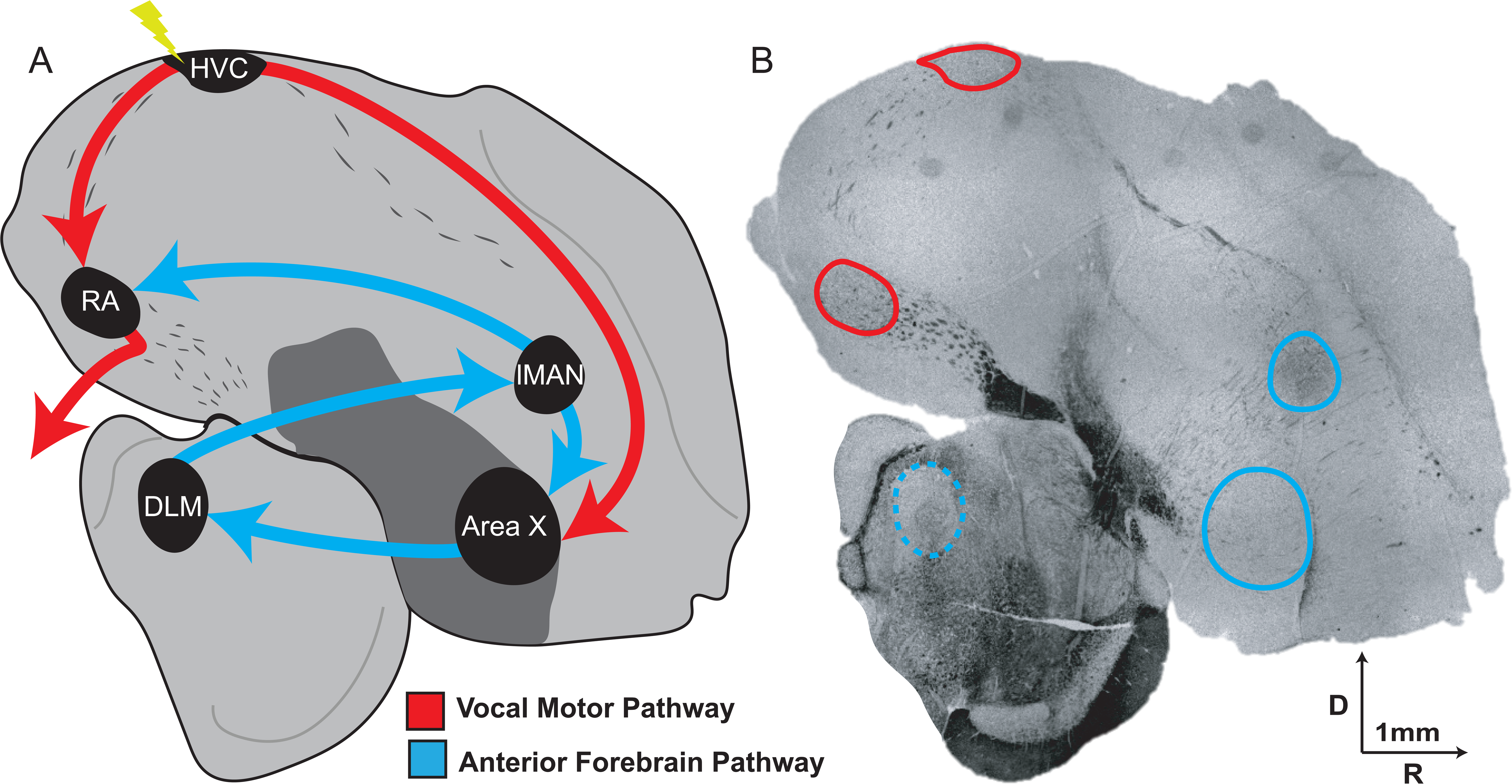
Schematic representation of selected zebra finch song nuclei and some interconnections. (A) Illustration of principal song system regions and pathways. Microlesions (yellow bolt) target the pre-motor cortical-like region HVC (proper name), which projects to the vocal motor nucleus RA (robust nucleus of the arcopallium; red arrows, vocal motor pathway) and the striatal/basal ganglia Area X (blue arrows, anterior forebrain pathway or AFP, a cortico-basal ganglia-thalamic loop critical for vocal learning and sensorimotor maintenance). Additional AFP components include lMAN (lateral magnocellular nucleus of the anterior nidopallium) and DLM (medial portion of the dorsolateral thalamic nucleus, dashed line indicates region is not present in the section shown). (B) Representative coronal section (darkfield image) used to identify song nuclei borders for lesioning, microdissection, and immunohistochemical analyses. Rostral is to the right, dorsal up; arrows = 1 mm. Note the distinct nuclear organization of song regions, contrasting with laminated mammalian cortex, facilitating targeted manipulations in this model.

**Figure 2:**
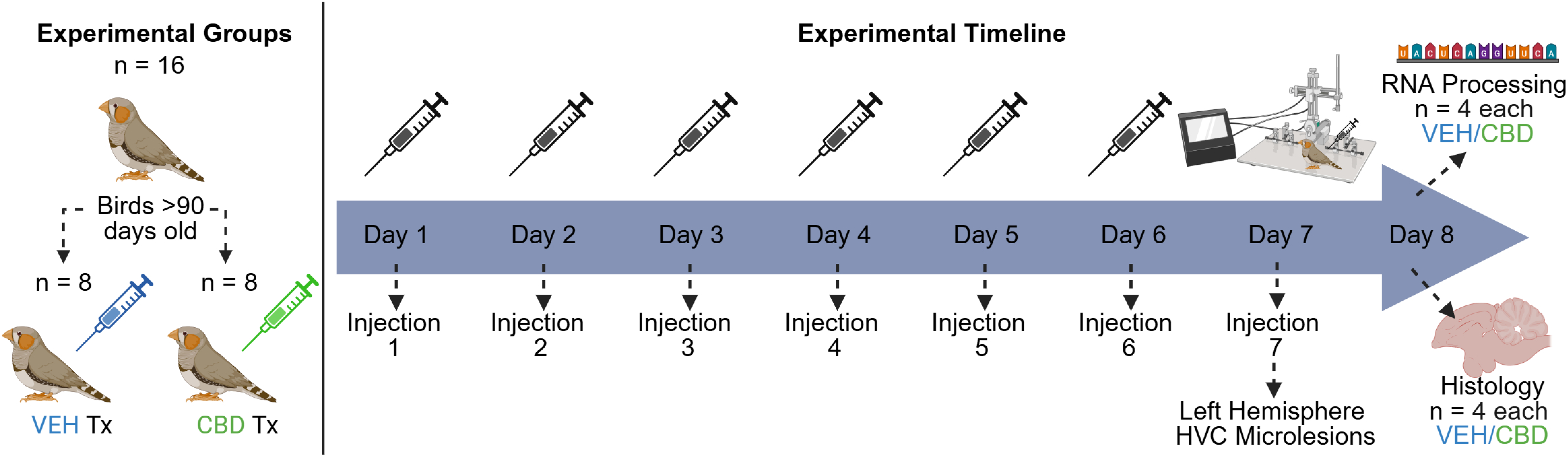
Experimental design and treatment timeline. Adult male zebra finches (> 90 days post-hatch; n = 16 total) were randomly assigned to vehicle (VEH; n = 8) or cannabidiol (CBD; n = 8) treatment groups. Animals received once-daily intramuscular injections (50 µl volume) for 6 consecutive days (Days 1–6), with CBD dosed at 10 mg/kg. This repeated pretreatment regimen was used to allow time for CBD accumulation in tissues (given its high lipophilicity and prolonged elimination half-life) to approach steady-state levels prior to surgery. On Day 7, a final pre-operative injection was administered, immediately followed by unilateral HVC microlesion surgery. Birds were euthanized 24 h post-surgery (Day 8) to capture acute injury responses. For histological analyses (n = 4 per group), birds were transcardially perfused with paraformaldehyde and brains fixed for sectioning. For RNA analyses (n = 4 per group), brains were rapidly extracted, and fresh tissue microdissected from song control nuclei (HVC, RA, Area X) for immediate RNA processing.

Partial HVC microlesions (∼10% ablation) disrupt song transiently, resolving in about 10 days via auditory-dependent adult relearning [16–20]. Unilateral lesions provide powerful within-subject controls while enabling examination of downstream effects in projection targets (robust nucleus of the arcopallium [RA] for motor execution and Area X for adaptive plasticity), including circuit-wide propagation of injury responses such as neuroinflammation, oxidative stress, and synaptic loss (e.g., Tripson et al., 2023). This TBI model thus offers a behaviorally quantifiable, mechanistically tractable system for investigating focal injury and recovery in a specialized circuit for sensorimotor learning and production.

Prior work demonstrated that daily CBD treatment (10 mg/kg) significantly reduces the magnitude of vocal disruption and accelerates recovery following HVC microlesions [16]. These behavioral improvements are mechanistically linked to robust anti-inflammatory effects (reduced pro-inflammatory cytokines IL6, IL1B, and TNFA; increased anti-inflammatory IL10), decreased oxidative stress (lower SOD2 expression and DHE fluorescence), attenuated microglial marker TMEM119, preserved synaptic densities (VGLUT2/PSD95 colocalization), NRF2 pathway activation (increased nuclear pNRF2), and upregulation of plasticity-related genes (BDNF, ARC/ARG3.1, MSK1) across song control nuclei [20]. This coordinated pattern of homeostatic and protective responses—spanning inflammation suppression, redox balance, synaptic stability, and plasticity promotion, strongly suggests involvement of additional glial cell types beyond microglia, such as astrocytes, which are known to orchestrate many of these processes in neural circuits under stress.

For example, synaptic stability post-injury requires coordinated neuron-glia interactions [21–23]. Astrocytes help maintain this stability by regulating redox balance, cytokine signaling, and neurotrophic support [24–26]. Within astrocytes responding to CNS injury, “reactivity programs” are triggered: coordinated changes in gene expression that drive morphological (e.g., process extension/retraction) and metabolic shifts. Neuroinflammatory signals from microglia (e.g., IL1B, TNFA, C1Q) can trigger neurotoxic phenotypes with upregulated complement components like C3 and pro-inflammatory genes [27]. In contrast, non-inflammatory insults (e.g., ischemia/hypoxia) often elicit a neuroprotective phenotype featuring growth factor release and production of repair molecules. However, astrocyte reactivity is highly heterogeneous and depends on the specific insult, with outcomes ranging from protective to detrimental. Recent consensus moves beyond rigid binary labels such as inflammatory/neurotoxic “A1-like” or anti-inflammatory/neuroprotective “A2-like”, instead recommending description by specific molecular and functional changes [28]. Astrocyte dysfunction generally worsens neuronal vulnerability and impairs recovery, while maintained homeostasis promotes circuit resilience [29]. It is worth noting that endocannabinoid modulation (e.g., CBD acting via PPARG/TRPV channels) is reported to enhance astrocyte-mediated recovery after traumatic brain injury [30].

The zebra finch model is particularly well-suited for studying region-selective glial responses in specialized vocal circuits that parallel those controlling human speech [9]. Focal brain injury induces robust astrogliosis, characterized by upregulation of glial androgen receptors and aromatase, which supports local estrogen synthesis and neuroprotection [31,32]. Vocal nuclei exhibit distinct patterns of astrocyte marker expression: GFAP-positive astrocytes are sparse and restricted (primarily to pallial borders, vasculature, and white matter, whereas GS-positive astrocytes (that we focus on here) are abundant and homogeneously distributed throughout the pallium, including HVC, RA, and Area X [33,34]. These differential morphologies and distributions likely contribute to circuit-specific glial functions.

We hypothesized that CBD modulates astrocyte responses to focal HVC injury in a region-selective manner, thereby stabilizing song circuits through distinct cellular mechanisms. The songbird model provides a unique opportunity to identify these mechanisms and their contributions to the selective recovery of specific sensorimotor circuit elements. Building on prior work showing that CBD can shift injury responses away from robust astrogliosis and neurotoxic inflammation toward homeostatic, protective phenotypes, we demonstrate here that CBD enhances post-lesion recovery by preserving astrocyte viability, attenuating injury-induced lysosomal stress and reactivity programs, and enhancing metabolic and antioxidant capacity across vocal-motor and vocal-learning circuit nodes (HVC, RA, and Area X). These glial outcomes parallel recent independent evidence that cannabidiol can engage TrkB signaling via direct interaction with the adaptor protein FRS2, leading to downstream attenuation of JAK2/STAT3-mediated neuroinflammation and preservation of synaptic integrity in chronic neurodegenerative models [35].

## Methods

### Materials

Unless stated otherwise, materials and reagents were obtained from MilliporeSigma or Thermo-Fisher Scientific. Cannabidiol (CBD) was supplied by GW Research Ltd. (Cambridge, UK) as a crystalline powder with >98% purity. CBD was suspended in vehicle to achieve a final dosing of 10 mg/kg in 50 μl. The vehicle consisted of a 2:1:17 ratio of ethanol:Alkamuls EL-620 (Rhodia, Cranbury, NJ):phosphate-buffered saline. The resulting ethanol dosage was 0.33 mg/kg per day, which is lower than levels voluntarily self-administered by zebra finches [36]. Isoflurane (Isospire, Dechra #17033-091-25), used for anesthesia, was provided by the Department of Comparative Medicine at East Carolina University.

### Drug treatments

CBD stock solutions, prepared as described above, were stored in sterile 5 ml septum-capped vials at 4 °C and refreshed at least once per week. For injections, drug solutions were loaded into sterile 1 cc insulin syringes fitted with 30-gauge needles. On injection days, birds were captured by hand in the early morning while lights remained off. The pectoralis muscle was exposed by parting the feathers with a small amount of 70% ethanol applied via squirt bottle. A 50 µL injection was administered into one of four quadrants of the pectoralis muscle, with the site rotated daily to reduce the risk of localized tissue damage from repeated dosing.

### Experimental timeline

Experiments were conducted over an 8-day period. Birds received six daily intramuscular injections of 50 μL CBD prior to surgery to allow the drug—given its lipophilic nature, large volume of distribution, and prolonged elimination half-life—to approach steady-state levels. On day 7, a final pre-operative injection was administered, followed by microlesion surgery as described below. Birds were euthanized 24 hours later (day 8). Brains were either perfused and fixed in paraformaldehyde or rapidly collected fresh for RNA isolation.

### Animals and environment

Adult male zebra finches (>= 90 days) were raised in the laboratory’s breeding aviary and maintained under standard conditions (78 °F; 12:12 h light/dark cycle). Only males were used in the study due to their capacity for song production. Birds were housed individually in standard finch cages (9″ × 11″ × 17″) with unrestricted access to food and water. While birds were visually isolated from conspecifics, they were not completely acoustically isolated. Sound-attenuated recording chambers made identification of recorded subjects unequivocal. All procedures were approved by the East Carolina University Animal Care and Use Committee. IF and qRT-PCR experiments were done using tissue collected from four adult zebra finches per treatment group. Original plans were to employ groups of eight divided into two cohorts of four birds each. Effect sizes for most measures were great enough that clear treatment differences were easily statistically distinguished after the first cohorts. In the interest of minimizing vertebrate animal impact, we did not add a second cohort as originally planned.

### HVC microlesion surgery

Unilateral microlesions of the HVC were performed using previously established methods. Lesions were targeted to the left hemisphere based on evidence of lateralization patterns in zebra finches that resemble those seen in human language, highlighting the functional similarities between birdsong and speech [37]. Furthermore, our prior work demonstrated that left-hemisphere HVC lesions result in more pronounced vocal impairments compared to right-hemisphere lesions [20]. Prior to surgery, birds received bupivacaine (0.05%, ∼75 µL) as a local anesthetic and were anesthetized with isoflurane. Birds were positioned in a stereotaxic apparatus with the bifurcation of the midsagittal sinus serving as the stereotaxic zero. Small craniotomies were made above the left HVC. To achieve partial (∼10%) HVC damage, two lesion sites were targeted: 2.4 mm and 2.8 mm lateral to stereotaxic zero, each to a depth of 0.6 mm. Lesions were induced by applying 100 μA of current for 35 s at each site using a PFA-coated tungsten electrode (A-M Systems; 0.008 in bare diameter, 0.011 in coated diameter), with the insulation removed at the electrode tip to expose the tungsten at the lesion site. Birds recovered in a warmed incubator before being returned to their recording chambers.

### Quantitative RT-PCR

As discussed above, groups of n = 4 birds were employed. cDNA samples were analyzed in triplicate (technical replicates). For each animal, 1 mm diameter biopsy punches were obtained from 2 mm thick parasagittal sections of the telencephalon (see Fig. 1) using RNase-free techniques (sterile, RNase-free gloves and materials). Punches were used to isolate tissue from three brain regions of interest—HVC, RA, and Area X—from both hemispheres. The unlesioned right hemisphere served as an internal control, while the lesioned left hemisphere represented the injury condition. Tissue samples were homogenized in TRIzol reagent (Invitrogen, 15,596,026), followed by phase separation with chloroform and RNA precipitation using isopropanol. The resulting RNA pellets were washed and resuspended in RNase-free water, and RNA concentration and purity were assessed using a NanoDrop spectrophotometer. Complementary DNA was synthesized from 250 ng of total RNA using an iScript cDNA synthesis kit (Bio-Rad, 1,708,890). Completed cDNA reactions were diluted five-fold using RNase-free water prior to quantitative PCR. Quantitative PCR was performed using SYBR Green Supermix (Bio-Rad, 1,725,271). Amplification specificity was verified by melt curve analysis, and cycle threshold (Ct) values were obtained using Design and Analysis Software version 2.8.0 (Thermo Fisher Scientific). Gene expression levels were normalized to the reference gene TBP, and relative expression was calculated using the ΔΔCt method with the unlesioned hemisphere as the comparator. Primer sequences are provided in Supplemental Table S1.

### Tissue harvesting and processing

Following euthanasia and transcardial perfusion with phosphate-buffered saline followed by 4% paraformaldehyde (PFA), brains were post-fixed overnight in 4% PFA at 4 °C. Tissue was then cryoprotected in 20% sucrose until brains sank, indicating sufficient water displacement and infiltration. Brains were bisected at the midline, embedded in OCT compound, and rapidly frozen using a slurry of dry ice and 2-methylbutane. Parasagittal sections (10 µm) were cut using a cryostat maintained at −20 °C. Sections from both hemispheres were mounted onto Superfrost Plus slides and stored at −20 °C until further processing.

### Immunofluorescent staining

Unless otherwise stated, all steps were performed at room temperature (25 °C). To reduce autofluorescence from lipofuscin, sections were treated with TrueBlack® Lipofuscin Autofluorescence Quencher (Cell Signaling Technology, Cat. #92401) according to the manufacturer’s instructions. Briefly, slides were incubated in TrueBlack working solution, rinsed thoroughly in phosphate-buffered saline (PBS), and allowed to equilibrate prior to downstream processing. Following autofluorescence quenching, sections were permeabilized in 0.2% Triton X-100 for 2 minutes and then blocked in 5% normal goat serum for 1 hour. Primary antibodies were diluted in 2% normal goat serum and applied at optimized concentrations, including anti-glutamine synthetase (1:500; Synaptic Systems, Cat. #367004), anti-LAMP1 (1:500; Abcam, ab24170), anti-cleaved caspase-3 (Asp175,1:500; Cell Signaling Technology, Cat. #9661), and anti-NeuN (clone A60,1:500; Sigma-Aldrich, Cat. #ZMS377). Primary antibodies were applied in separate staining panels or in compatible combinations, depending on host species and fluorophore compatibility. Slides were incubated with primary antibodies overnight at 4 °C. The next day, slides were incubated with secondary antibodies—Alexa Fluor 647 goat anti-mouse IgG (H+L) (Thermo Fisher Scientific Cat. #A-21235 1:500) Alexa Fluor 555 goat anti-guinea pig IgG (H+L) (Thermo Fisher Scientific Cat. #A-21435 1:500) and Alexa Fluor 488 goat anti-rabbit IgG (H+L) (Thermo Fisher Scientific Cat. #A-11008 1:500)—diluted in 2% normal goat serum for 1 hour. After secondary incubation, slides were washed twice with PBS (5 minutes each), counterstained with Hoechst (1:10,000) and washed a final time. Coverslips were mounted using Fluoro-Gel with Tris Buffer mounting medium (Electron Microscopy Sciences).

### Fluorescent imaging

Regions of interest included HVC (excluding infarcted areas), RA, and Area X. To ensure consistent anatomical representation across treatment conditions, sections containing portions of all three regions were selected for imaging. Fluorescent images were acquired using a Keyence BZ-X800 fluorescence microscope equipped with a 20× Plan Apo objective (NA 0.75; Keyence, model BZ-PA20). Images were captured using DAPI, GFP, TXRED, and Cy5 filter sets. For each region, three non-overlapping images were collected to ensure representative sampling across the area and to minimize localized variability. Image thresholding parameters were held constant across all images to ensure uniform quantification.

### Image analysis

Image analysis was performed using ImageJ software (NIH). Fluorescent images were converted to 8-bit grayscale and thresholded using consistent parameters across all animals and conditions. For cell identification and cleaved caspase-3 (CC3) analysis, Hoechst-stained nuclei were thresholded using the default method, segmented using the watershed function, and subjected to particle analysis with a size constraint of 10–300 µm² to identify individual nuclei. Nuclear masks were applied to additional channels, and logical AND operations were used to assess marker overlap. Nuclei overlapping NeuN or glutamine synthetase (GS) signal were classified as neuronal or astrocytic, respectively, and nuclei overlapping CC3 signal were classified as apoptotic. Cell counts were expressed as the percentage of total Hoechst-positive cells for each cell type (neurons and astrocytes), and CC3-positive cells were expressed as the percentage of the corresponding cell type. For lysosomal analysis, the Triangle thresholding method was applied to the GS channel to identify astrocytes, which were then segmented using the watershed function and analyzed with a size constraint of 50–600 µm². Astrocyte-derived masks were used to define astrocyte-specific regions of interest, within which a second particle analysis was performed on the LAMP1 channel (size constraint: 0–30 µm²). Lysosomal measurements were expressed as average lysosome size (µm²).

### Statistical analysis

Statistical analyses were performed using GraphPad Prism version 11.0.0. Immunofluorescence (IF) data were analyzed using three-way mixed-effects models using restricted maximum likelihood (REML) estimation. Models included Bird ID as a random factor, Region (HVC, RA, Area X) × Lesion Condition (lesioned hemisphere or un-lesioned internal control) × Treatment (Vehicle or CBD), with dependent variables within Bird ID and across region treated as repeated measures where appropriate. When at least one interaction involving Region was statistically significant, data were stratified by region and post hoc analyses were performed using two-way ANOVA (Lesion Condition × Treatment) within each region, with multiple comparisons corrected using Tukey’s test. Quantitative RT-PCR gene expression data were analyzed using two-way mixed-effects models (Region × Treatment), with region treated as a repeated-measures factor within subjects, followed by Sidak’s multiple comparisons correction. IF data are presented as estimated means ± standard deviation (SD), while qRT-PCR data are presented as estimated means ± standard error of the mean (SEM). A p-value ≤ 0.05 was considered statistically significant. Complete statistics, including fixed effects from mixed-effects models (Table S1) and post hoc multiple comparisons (Table S2), are provided in the Supplementary Material.

## Results

### CBD increases astrocyte percentages in a region-dependent manner

We first examined whether cannabidiol (CBD) alters astrocyte populations across song nuclei, given the central role of astrocytes in coordinating inflammatory and metabolic responses following injury (Fig. 3D). A three-way mixed-effects model revealed significant main effects of region (p < 0.0001) and treatment (p = 0.0004), and interactions including Region × Lesion condition (p = 0.0039) and Region × Treatment (p = 0.006) for astrocytes as a percentage of total cells (i.e., Hoechst-labeled cell nuclei, see supplementary Table S1 for complete three-way model statistics and Table S2 for complete post hoc comparisons. The Region × Lesion interaction indicates that injury effects vary across song nuclei, consistent with region-specific susceptibility to secondary injury processes. The Region × Treatment interaction indicates that CBD effects are not uniform across song nuclei, suggesting region-dependent modulation of astrocyte responses. The Lesion × Treatment interaction indicates that CBD effects depend on injury status, supporting an injury-dependent mechanism by which CBD preferentially modulates astrocyte responses triggered by lesioning of tissue. Consistent with these interactions, post hoc comparisons revealed that CBD increased the percentage of GS+/Hoechst+ nuclei on the lesioned side in HVC (VEH: 12.92% vs CBD: 18.25%, 95% CI [−7.85%, −2.81%], p = 0.0003), across both hemispheres in RA (lesioned: VEH: 12.16% vs CBD: 19.96%, 95% CI [−8.17%, −0.51%], p = 0.0036; unlesioned: VEH: 11.83% vs CBD: 21.45%, 95% CI [−14.60%, −4.64%], p = 0.0007), and on the lesioned side in Area X (VEH: 6.34% vs CBD: 10.68%, 95% CI [−12.78%, −2.81%], p = 0.0269), with no change on the unlesioned side in Area X (p > 0.05). Together, these findings indicate that CBD drives region- and injury-dependent increases in astrocyte representation, supporting a role for astrocytes as key mediators of CBD-modulated responses to neural injury.

**Figure 3:**
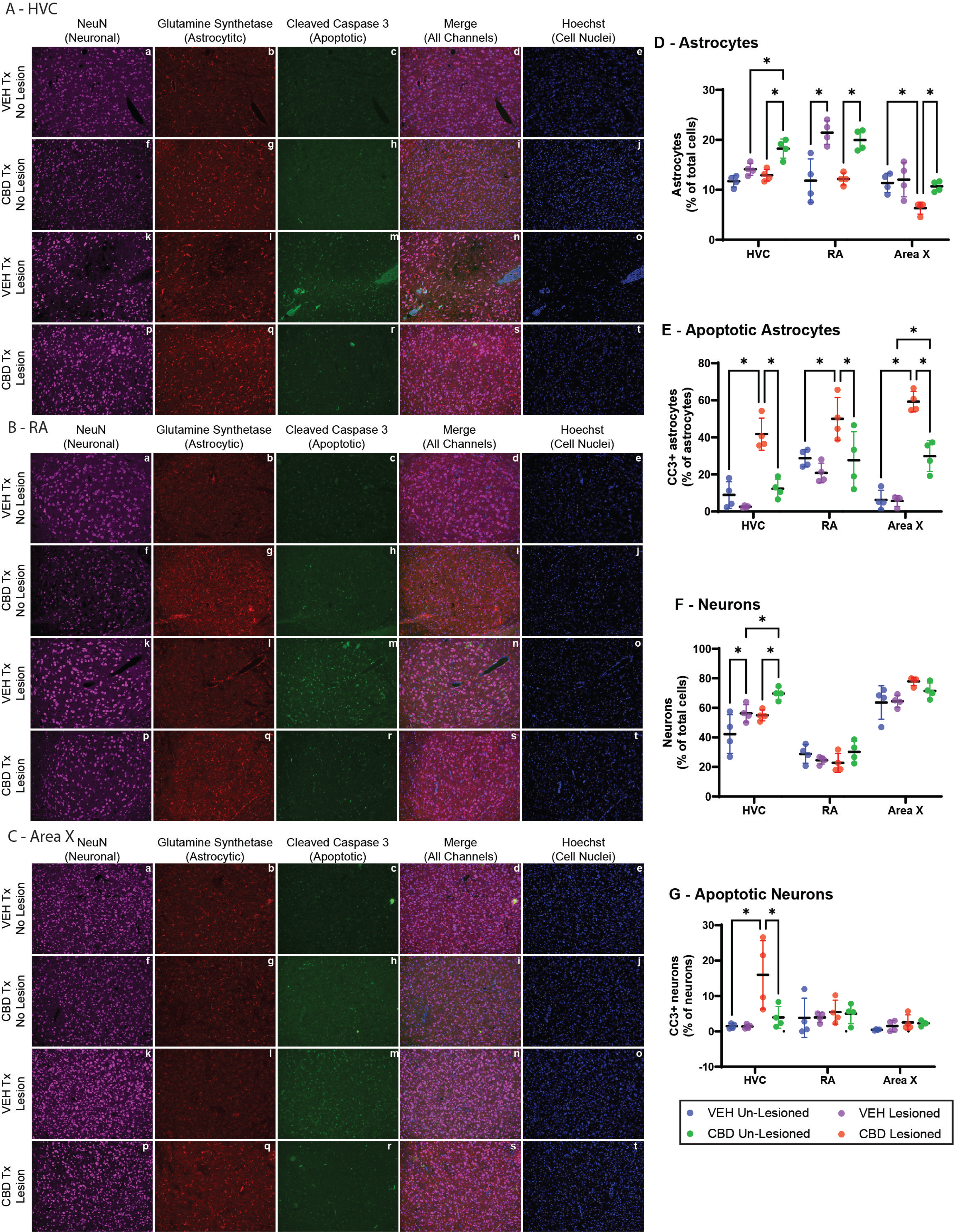
Cannabidiol (CBD) preserves astrocyte viability and reduces lesion-induced apoptosis in song control nuclei following HVC microlesions. Representative immunofluorescence images (A–C) and quantitative analyses (D–G) of NeuN (neuronal marker, magenta), glutamine synthetase (GS; astrocytic marker, red), cleaved caspase-3 (CC3; apoptotic marker, green), merged channels, and Hoechst nuclear counterstain (blue) in HVC (A), RA (B), and Area X (C) from vehicle (VEH)- and CBD-treated zebra finches. Scale bars = 200 µm. Quantitative data (D–G) depict cell populations expressed as: (D) GS+ astrocytes as a percentage of Hoechst+ cells: (E) CC3+ apoptotic cells as a percentage of Hoechst+ cells; (F) NeuN+ neurons as a percentage of Hoechst-positive cells. (D) Astrocytes (GS⁺) are significantly preserved in CBD-treated animals across HVC, RA, and Area X following lesions. (E) Apoptotic astrocytes (GS⁺/CC3⁺) are markedly reduced by CBD treatment in all three brain regions. (F) Neuronal density (NeuN⁺) shows modest preservation in HVC with CBD. (G) Apoptotic neurons (NeuN⁺/CC3⁺) remain low and largely unaffected by lesion or treatment. Data points represent individual animals (n = 4 per group. Statistical comparisons were performed using mixed-effects ANOVA with Sidak’s post-hoc test (*p < 0.05). These results demonstrate that CBD-mediated neuroprotection following focal HVC injury primarily involves enhanced astrocyte survival and suppression of apoptotic signaling in astrocytes, with secondary benefits to neuronal populations.

### CBD attenuates lesion-induced astrocyte apoptosis across song nuclei

To determine whether these shifts reflect altered survival, we next assessed astrocyte apoptosis, defined as the percentage of astrocytes positive for cleaved caspase-3 (CC3; CC3+ astrocytes out of total astrocytes identified by GS- and Hoechst-double labeling, see Fig. 3E). A three-way mixed-effects model revealed significant main effects of region (p = 0.0005), lesion condition (p < 0.0001), and treatment (p = 0.0008), together with Region × Lesion condition (p = 0.0007) and Lesion condition × Treatment (p = 0.0027) interactions for astrocyte apoptosis (three-way model: Table S1; post hoc comparisons: Table S2). The Region × Lesion interaction indicates that injury-induced astrocyte apoptosis varies across song nuclei. The Lesion × Treatment interaction indicates that CBD effects depend on injury status, consistent with selective modulation of astrocyte survival in lesioned tissue. Consistent with these interactions, post hoc comparisons revealed that lesions increased percentages of apoptotic astrocytes in HVC (VEH: 8.95% un-lesioned hemispheres vs 41.76% within lesioned hemispheres, 95% CI [−43.26%, −22.37%], p = 0.0002), RA (VEH: 28.75% vs 50.00%, 95% CI [−42.57%, 0.065%], p = 0.0498), and Area X (VEH: 6.20% vs 59.31%, 95% CI [−63.75%, −42.47%], p < 0.0001). CBD reduced % apoptotic astrocytes on the lesioned side in HVC (VEH: 41.76% vs CBD: 12.32%, 95% CI [18.27%, 40.63%], p < 0.0001), RA (VEH: 50.00% vs CBD: 27.72%, 95% CI [3.80%, 40.76%], p = 0.019), and Area X (VEH: 59.31% vs CBD: 29.92%, 95% CI [18.83, 39.96], p < 0.0001). Together, these findings indicate that lesions robustly increase astrocyte apoptosis across song nuclei, whereas CBD attenuates this response in an injury-dependent manner.

### CBD selectively increases percentages of neurons within HVC

Given the role of astrocytes in supporting neuronal maintenance within song nuclei (Fig. 3F), we next assessed whether astrocyte-associated changes were accompanied by altered percentages of neurons, defined as NeuN+ cells / total Hoechst+ cells × 100%. A three-way mixed-effects model revealed significant main effects of region (p < 0.0001) and lesion condition (p = 0.0075), along with Region × Lesion condition (p = 0.0116) and Region × Treatment (p = 0.0022) interactions for neuronal percentages (three-way model: Table S1; post hoc comparisons: Table S2). The Region × Lesion interaction indicates that injury-related changes in neuronal percentages vary across song nuclei. The Region × Treatment interaction indicates that CBD effects are region-specific, suggesting selective modulation of neuronal populations. Consistent with these interactions, post hoc comparisons revealed that CBD increased neuronal percentages in HVC on both the un-lesioned side (VEH: 42.25% vs CBD: 56.27%, 95% CI [−27.98%, −0.07%], p = 0.0489) and lesioned side (VEH: 54.93% vs CBD: 69.72%, 95% CI [−28.74%, −0.83%], p = 0.0378), with no effect detected in RA or Area X (p > 0.05). These findings indicate that CBD selectively increases neuronal percentages in HVC in a region-specific manner.

### CBD reduces lesion-associated neuronal apoptosis in HVC

We then examined neuronal apoptosis, expressed as the percentage of neurons positive for cleaved caspase-3 (CC3; CC3+ neurons / total neurons identified by NeuN and Hoechst double-labeling), to determine whether injury and treatment effects on neurons paralleled those observed in astrocytes (Fig. 3G). A three-way mixed-effects model revealed significant main effects of region (p = 0.0177) and lesion condition (p = 0.0267), together with Region × Lesion condition (p = 0.0039), Region × Treatment (p = 0.0379), and Region × Lesion condition × Treatment (p = 0.0212) interactions for neuronal apoptosis (three-way model: Table S1; post hoc comparisons: Table S2). The Region × Lesion interaction indicates that injury-induced neuronal apoptosis varies across song nuclei. The Region × Treatment and three-way interactions indicate that CBD effects on neuronal apoptosis are region- and injury-dependent. Consistent with these interactions, post hoc comparisons revealed that lesion increased percentages of apoptotic neurons in HVC (VEH: 1.47% vs 15.99%, 95% CI [−25.04%, −3.984%], p = 0.013), and that CBD reduced percentages of apoptotic neurons on the lesioned side in HVC (VEH: 15.99% vs CBD: 3.96%, 95% CI [2.836%, 21.22%], p = 0.0117), with no effects detected in RA or Area X (p > 0.05). These findings indicate that lesion-induced percentages of apoptotic neurons is highest in HVC and selectively attenuated by CBD in this region.

### CBD attenuates lysosomal area following injury

Given the consistent astrocyte-associated effects, we next examined lysosomal structure, as indicated by lysosome area (µm²) reflects cellular stress (Fig. 4). A three-way mixed-effects model revealed a significant Region × Lesion condition × Treatment interaction (p = 0.017; Table S1). This interaction indicates that CBD effects on lysosome size are both region- and injury-dependent. Consistent with this, post hoc comparisons revealed that CBD reduced lysosome area in HVC within both the un-lesioned side (VEH: 11.22 µm² vs CBD: 6.36 µm², 95% CI [2.41 µm², 7.31 µm²], p = 0.0006) and lesioned side (VEH: 11.53 µm² vs CBD: 8.17 µm², 95% CI [0.91 µm², 5.81 µm²], p = 0.0088), in RA on the un-lesioned side (VEH: 9.44 µm² vs CBD: 8.49 µm², 95% CI [2.649 µm², 7.297 µm²], p = 0.0003) but not the lesioned side (p > 0.05), and in Area X on both the un-lesioned side (VEH: 8.78 µm² vs CBD: 5.73 µm², 95% CI [1.07 µm², 5.03 µm²], p = 0.0039) and lesioned side (VEH: 10.28 µm² vs CBD: 6.67 µm², 95% CI [1.64 µm², 5.59 µm²], p = 0.0011). Together, these findings indicate that CBD broadly attenuates lysosomal area across song nuclei, with a reduced effect in the lesioned RA.

**Figure 4:**
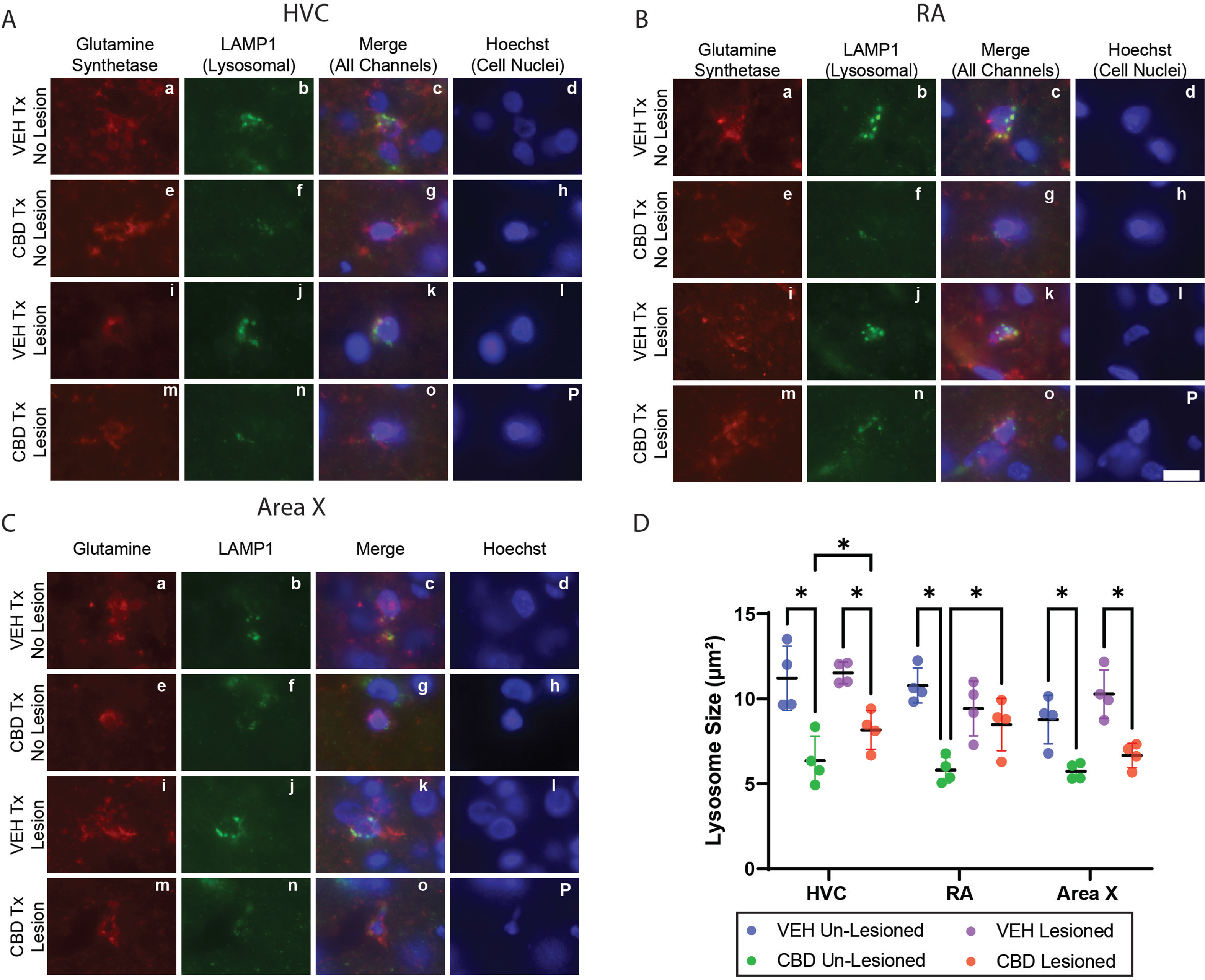
Cannabidiol (CBD) reduces astrocytic lysosomal size in song control nuclei following injury. Representative immunofluorescence images from HVC (A), RA (B), and Area X (C) in un-lesioned and lesioned hemispheres of vehicle (VEH)- and CBD-treated birds. Astrocytes were labeled with glutamine synthetase (GS; red), lysosomes with LAMP1 (green), and nuclei with Hoechst (blue); merged channels are shown. (D) Quantification of mean astrocytic lysosomal area (µm²; LAMP1+ puncta per GS+ cell) reveals that CBD treatment significantly reduces lysosomal size bilaterally across HVC, RA, and Area X compared to VEH controls (p < 0.05). In the lesioned hemisphere of RA, CBD-treated animals showed a trend toward reduction relative to VEH but did not reach statistical significance. n = 4 birds per group. Statistical comparisons were performed using mixed-effects ANOVA with Sidak’s multiple comparisons test. Bar = 10 microns

### CBD reduces lysosomal and autophagy-associated gene expression

To determine whether these lysosomal area changes were accompanied by transcriptional responses, we next quantified mRNA expression using qRT-PCR, expressed as fold change relative to each animal’s contralateral control hemisphere (Fig. 5). A two-way mixed-effects model revealed significant main effects of region and treatment, as well as a Region × Treatment interaction for LAMP1 mRNA expression (two-way model: Table S1; post hoc comparisons: Table S2). The Region × Treatment interaction indicates that CBD effects on lysosomal gene expression are region-dependent. Consistent with this, post hoc comparisons revealed that CBD reduced LAMP1 mRNA fold-expression in HVC (VEH: 2.42-fold vs CBD: 1.70 -fold, 95% CI [0.415-fold, 1.032-fold], p < 0.0001) and RA (VEH: 1.14-fold vs CBD: 0.53-fold, 95% CI [0.30-fold, 0.91-fold], p = 0.0002), with no effect in Area X (p > 0.05). A similar model for LC3 mRNA expression revealed significant effects of region (p < 0.0001), treatment (p = 0.0047), and their interaction (p < 0.0001; Fig. 5B). The Region × Treatment interaction indicates that CBD effects on autophagy-related gene expression are also region-dependent. Post hoc comparisons revealed that CBD reduced LC3 mRNA expression in HVC (VEH: 2.020-fold vs CBD: 1.394-fold, 95% CI [0.35-fold, 0.90-fold], p < 0.0001) and RA (VEH: 0.84-fold vs CBD: 0.45-fold, 95% CI [0.11-fold, 0.67-fold], p = 0.0048), while increasing expression in Area X (VEH: 0.59-fold vs CBD: 1.02-fold, 95% CI [−0.71-fold, −0.15-fold], p = 0.0022). Together, these findings indicate that CBD modulates lysosomal and autophagy-related transcriptional responses in a region-dependent manner.

**Figure 5:**
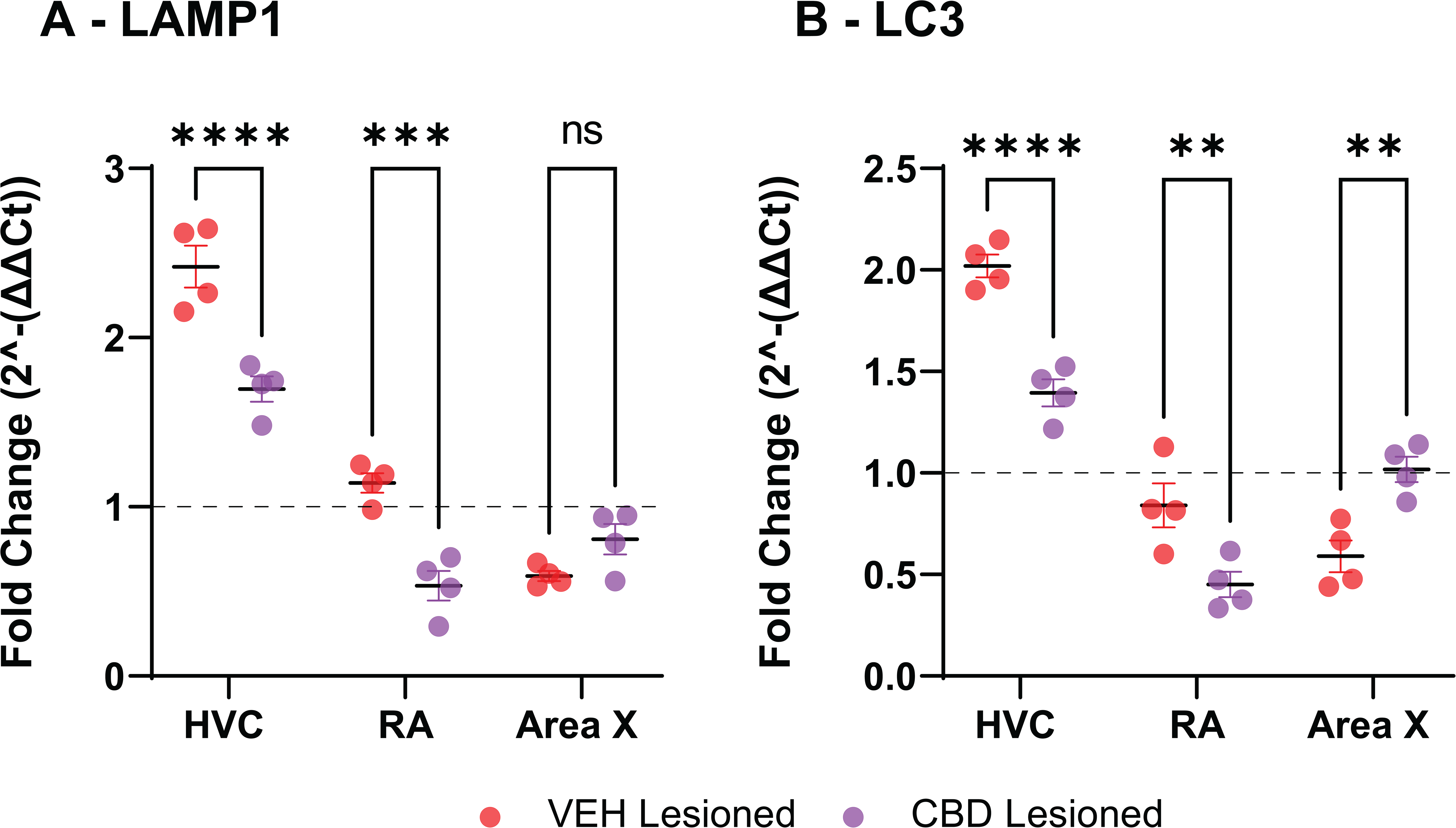
Cannabidiol (CBD) modulates expression of lysosomal and autophagosomal markers in song control nuclei post-injury. Quantitative RT-PCR analysis of lysosome- and autophagosome-related transcripts in lesioned hemispheres of HVC, RA, and Area X from vehicle- and CBD-treated birds, normalized to the contralateral un-lesioned hemisphere. (A) LAMP1 (lysosomal membrane marker) mRNA is significantly reduced in HVC and RA with CBD treatment, with no significant change in Area X. (B) LC3 (autophagosome marker) mRNA is significantly reduced in HVC and RA with CBD treatment but significantly increased in Area X. Data represent mean fold change ± SEM; individual points show n = 4 animals per group. Statistical comparisons: mixed-effects ANOVA with Sidak’s multiple comparisons test (p < 0.05).

### CBD enhances astrocyte metabolic and antioxidant gene expression

We next examined levels of astrocyte-associated metabolic and antioxidant transcripts, given the role of astrocytes in astrocyte-mediated glutamate clearance and redox regulation following injury (Fig. 6). mRNA expression was quantified using qRT-PCR and expressed as fold change relative to each animal’s contralateral control hemisphere. A two-way mixed-effects model revealed significant main effects of region (p = 0.0007) and treatment (p < 0.0001), along with a Region × Treatment interaction for glutamine synthetase (GS) mRNA (two-way model: Table S1; post hoc comparisons: Table S2). The Region × Treatment interaction indicates that CBD effects on astrocyte-associated metabolic gene expression are region-dependent. Consistent with this, post hoc comparisons revealed that CBD increased GS mRNA fold expression in HVC (VEH: 0.82-fold vs CBD: 1.85-fold, 95% CI [−1.51-fold, −0.57-fold], p < 0.0001) and RA (VEH: 0.43-fold vs CBD: 1.76-fold, 95% CI [−1.80-fold, −0.86-fold], p < 0.0001), with no effect in Area X (p > 0.05). Similarly, a model for GCLM mRNA revealed significant effects of region (p = 0.0001), treatment (p < 0.0001), and their interaction (p < 0.0001; Fig. 6B). The Region × Treatment interaction indicates that CBD effects on astrocyte-associated antioxidant gene expression are also region-dependent. Post hoc comparisons revealed that CBD increased GCLM mRNA fold expression in HVC (VEH: 0.66-fold vs CBD: 1.60-fold, 95% CI [−1.27-fold, −0.62-fold], p < 0.0001) and RA (VEH: 1.06-fold vs CBD: 1.44-fold, 95% CI [−0.71-fold, −0.06-fold], p = 0.018), with no effect in Area X (p > 0.05). These findings indicate that CBD enhances astrocyte-mediated glutamate handling and antioxidant capacity in a region-dependent manner following injury.

**Figure 6:**
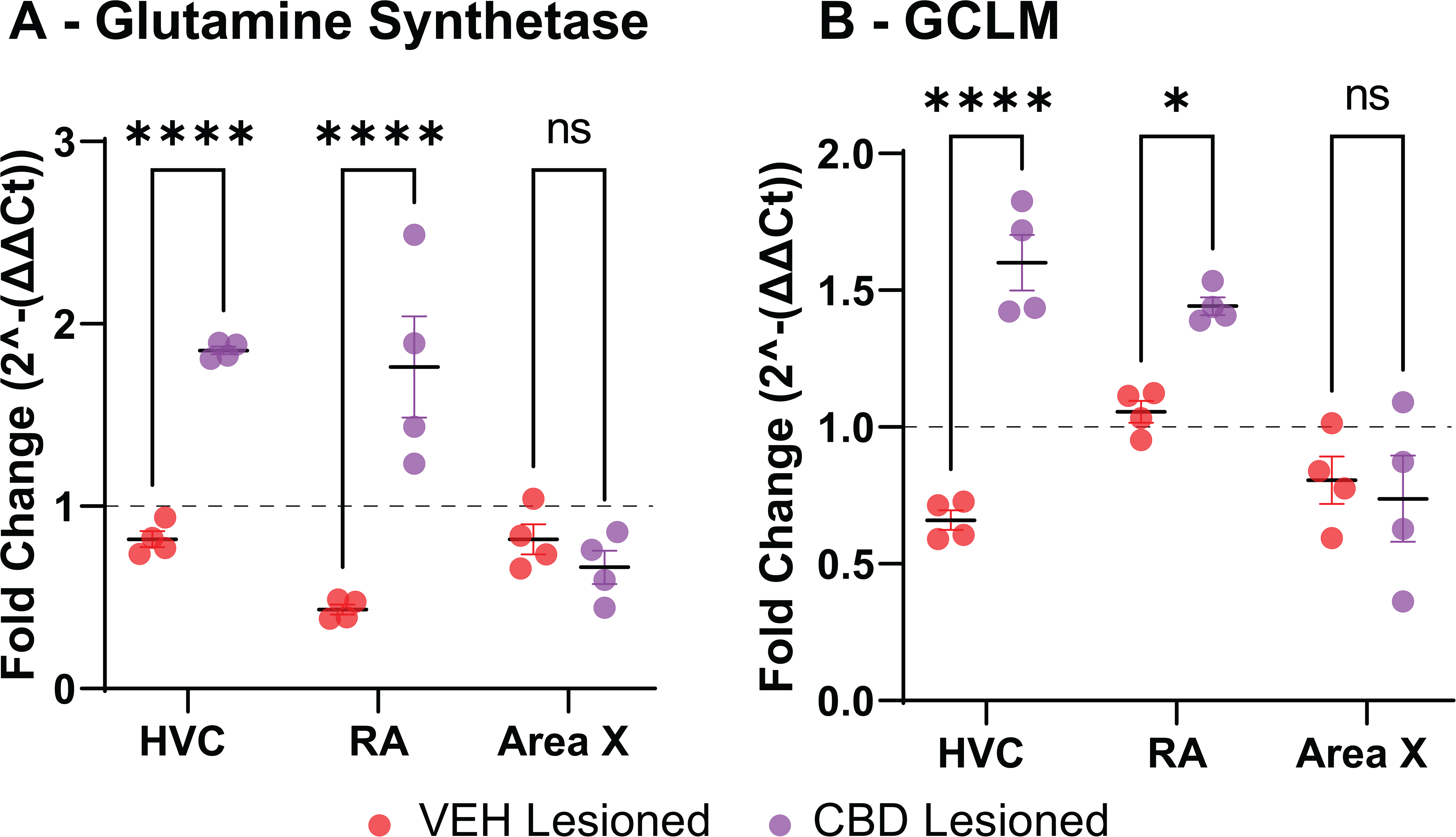
Cannabidiol (CBD) enhances expression of astrocytic metabolic and antioxidant marker transcripts in song control nuclei following injury. Quantitative RT-PCR analysis of selected transcripts in lesioned hemispheres of HVC, RA, and Area X from vehicle- and CBD-treated birds, normalized to the contralateral un-lesioned hemisphere. (A) Glutamine synthetase (GS), a key astrocytic enzyme supporting glutamate recycling and metabolic homeostasis, shows significantly increased mRNA levels in HVC and RA with CBD treatment. (B) Glutamate-cysteine ligase modifier subunit (GCLM), a regulator of glutathione synthesis and cellular antioxidant capacity, shows significantly increased mRNA levels in HVC and RA with CBD treatment. Data represent mean fold change ± SEM from n = 4 animals per group (individual points shown). Comparisons were performed using mixed-effects ANOVA with Sidak’s multiple comparisons test (p < 0.05).

### CBD dampens astrocyte reactivity-associated transcriptional responses

Taken together, the combined changes in astrocyte percentages, diminished astrocyte apoptosis, normalization of lysosomal structure, and restoration of metabolic gene expression point toward coordinated modulation of astrocyte activation state. We therefore assessed astrocyte reactivity-associated transcripts using qRT-PCR, expressed as fold change relative to each animal’s contralateral control hemisphere (Fig. 7). A two-way mixed-effects model revealed a Region × Treatment interaction for S100A10 mRNA (p = 0.0032; Fig. 7A), indicating region-specific effects of CBD on astrocyte reactivity. Consistent with this, post hoc comparisons revealed that CBD reduced S100A10 mRNA fold expression in HVC (VEH: 2.73-fold vs CBD: 1.51-fold, 95% CI [0.56-fold, 1.88-fold], p = 0.0004), with no effect in RA or Area X (p > 0.05). For C3 mRNA fold expression, significant main effects of region (p < 0.0001) and treatment (p = 0.0086) were observed (Fig. 7B). Post hoc comparisons revealed that CBD reduced C3 mRNA expression in HVC (VEH: 5.51-fold vs CBD: 3.96-fold, 95% CI [0.29-fold, 2.81-fold], p = 0.0136) and RA (VEH: 2.39-fold vs CBD: 0.93-fold, 95% CI [0.20-fold, 2.72-fold], p = 0.0207), with no effect in Area X (p > 0.05). Aromatase mRNA (a marker of astrocyte reactivity) showed significant effects of region (p < 0.0001), treatment (p = 0.0025), and their interaction (p < 0.0001; Fig. 7C), indicating region-dependent effects of CBD. Post hoc comparisons revealed that CBD reduced aromatase mRNA fold expression in HVC (VEH: 3.28-fold vs CBD: 1.01-fold, 95% CI [1.70-fold, 2.84-fold], p < 0.0001), with no effect in RA or Area X (p > 0.05). Collectively, these findings indicate that CBD dampens astrocyte reactivity-associated gene expression, with effects most pronounced in regions exhibiting injury-associated activation.

**Figure 7:**
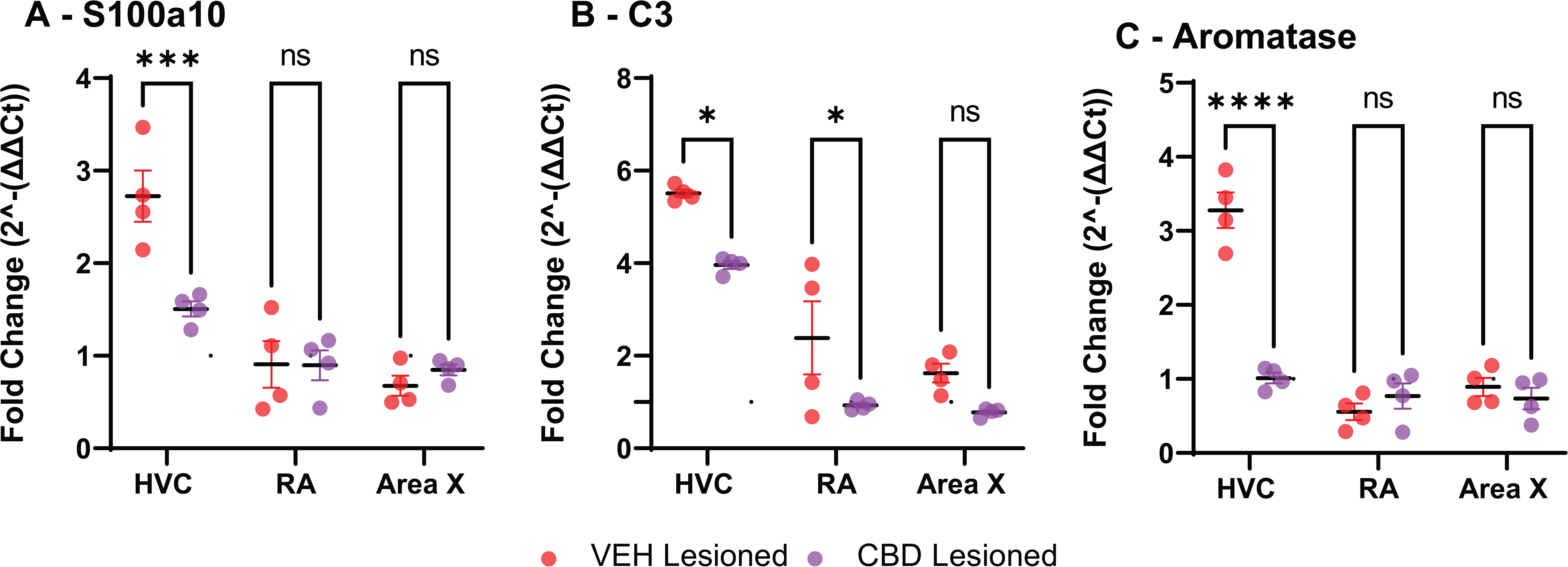
Cannabidiol (CBD) attenuates injury-induced astrocyte reactivity markers in song control nuclei. Quantitative RT-qPCR analysis of selected astrocyte-related transcripts in the lesioned hemispheres of HVC, RA, and Area X from vehicle- and CBD-treated zebra finches, expressed as fold change relative to the contralateral un-lesioned hemisphere (normalized to endogenous control). (A) S100A10 (marker of anti-inflammatory/pan-reactive astrocytes) is significantly reduced in HVC by CBD treatment, with no significant change in the projection targets RA and Area X. (B) C3 (marker associated with neurotoxic “A1-like” astrocyte reactivity) is significantly decreased by CBD in both HVC and RA. (C) Aromatase (injury-responsive astrocyte marker linked to local estrogen synthesis) is significantly reduced in HVC following CBD treatment. Data are presented as mean fold change ± SEM; individual points represent biologically independent animals (n = 4 per group). Statistical comparisons were performed using mixed-effects ANOVA followed by Sidak’s multiple comparisons test (p < 0.05, *p < 0.01, **p < 0.001; non-significant differences are not annotated). These findings are consistent with CBD-mediated suppression of lesion-induced astrocyte stress responses, complementing the preservation of astrocyte viability (Fig 3) and reduction in neuroinflammatory signaling reported previously.

## Discussion

This study directly builds on our earlier work demonstrating that cannabidiol (CBD, 10 mg/kg daily) protects vocal quality and accelerates recovery of learned song patterns after HVC microlesions in adult male zebra finches [16]. Vocal recovery was associated with robust CBD suppression of microlesion-induced neuroinflammation (reduced IL6, IL1B, and TNFA), oxidative stress (reduced SOD2 and DHE fluorescence), microglial activation (reduced TMEM119), and circuit-wide synaptic loss (preserved VGLUT2/PSD95 colocalization), as well as activation of NRF2 nuclear translocation with upregulation of plasticity-associated transcripts (BDNF, ARC/ARG3.1, MSK1, Tripson et al., 2023). We designed the present experiments to identify cellular substrates underlying these protective effects. We now show that astrocytes are a primary, though not exclusive, target of CBD-mediated neuroprotection.

These findings complement recent independent evidence that CBD interacts with the TrkB adaptor FRS2 to stabilize receptor signaling and inhibit JAK2/STAT3/SOCS1-driven inflammatory cascades [35]. In that work, FRS2-dependent TrkB activation was associated with reduced astrogliosis (GFAP), cytokine production, and synaptic loss: outcomes that align closely with the astrocyte-preserving effects, lowered reactivity markers (C3, S100A10, aromatase), and synaptic protection we observe across song-control nuclei after acute TBI. The present astrocyte-centric data therefore extend the known neuroprotective profile of cannabidiol to a vocal-learning model of focal injury, reinforcing the potential for glial-targeted mechanisms in circuit resilience.

CBD preserved astrocyte viability across regions of both the anterior forebrain pathway and vocal motor circuits (HVC, RA, and Area X; see Fig. 1), attenuated injury-induced lysosomal stress and reactivity programs, and enhanced mediators of astrocyte metabolic and antioxidant capacity. These glial-associated actions provide a mechanistic link between previously reported anti-inflammatory, anti-oxidative, and synaptoprotective outcomes and the accelerated behavioral recovery of song, a complex sensorimotor skill.

A central finding is that HVC microlesions trigger substantial astrocyte apoptosis, accounting for a large fraction of cleaved caspase-3–positive cells in the three song nuclei investigated (HVC, RA, and Area X; Fig. 3). CBD markedly reduces this astrocyte loss while preserving or expanding astrocyte populations in a region-dependent manner (Fig. 3D,E). This astrocyte-specific protection is particularly relevant to secondary injury processes, as the primary electrolytic damage remains constant across treatment groups while CBD limits the subsequent apoptotic, lysosomal, and reactivity cascades that would otherwise amplify circuit-wide dysfunction (see Alalawi et al., 2019; Tripson et al., 2023, for propagation of inflammatory and synaptic responses). This general, circuit-wide astrocytic resilience contrasts with more localized neuronal preservation (confined largely to the directly-lesioned HVC; Fig. 4) and aligns with evidence that astrocyte viability is a critical determinant of circuit recovery after focal CNS injury [38–40]. Astrocytes maintain neuronal survival and synaptic stability through metabolic coupling (e.g., lactate shuttling, glutamate homeostasis, redox buffering) and structural support [41,42]. Astrocytic aerobic glycolysis generates energy- and survival-promoting lactate that is shuttled to neurons via selective transporters, providing oxidative substrate while preserving neuronal redox balance and supporting synaptic plasticity [43]. Astrocytes also clear excess glutamate to protect against excitotoxicity and maintain synaptic homeostasis [44] and serve as the primary CNS source of glutathione for redox buffering and neuroprotection [45].

The endocannabinoid system (ECS) regulates many of these functions, and astrocytes express both canonical (CB₁, inducibly CB₂) and non-canonical ECS-linked receptors (TRPV1/4, PPARG; [25,46]). Although CBD is not a direct CB₁/CB₂ agonist, it modulates astrocytic signaling via PPARG and TRPV channels [47,48] consistent with the broad circuit-wide preservation of astrocyte populations observed here. By stabilizing astrocytes, CBD may prevent secondary propagation of injury signals that would otherwise amplify neuronal vulnerability and synaptic destabilization: deficits reversed in our prior study [20].

Injury also imposes significant lysosomal stress on astrocytes, as evidenced by enlarged LAMP1-immunopositive lysosomes and upregulation of LAMP1 and LC3 mRNA (Fig. 5). In the case of enlarged lysosomes, these were within GS-expressing astrocytes. In the case of LC3 upregulation, this may have also included extra-astrocytic expression (e.g. from microglia and or neurons). CBD normalized lysosomal morphology and suppressed these transcriptional responses, indicating reduced proteostatic stress rather than global lysosomal suppression. Lysosomal/autophagic pathways are tightly coupled to cellular redox balance and mitochondrial quality control [49]. Their dysregulation exacerbates oxidative stress and inflammation [50].

Reductions in lysosomal burden and LAMP1/LC3 expression are consistent with the hypothesis that CBD improves autophagic clearance of injury-related debris, thereby limiting accumulation of damaged organelles and proteins and promoting astrocyte and circuit resilience. However, the observed decreases could alternatively reflect reduced production of cellular debris upstream of autophagy (e.g., via CBD’s established suppression of neuroinflammation, oxidative stress, microglial activation, and synaptic loss; Tripson et al., 2023).

Pharmacological assessment of autophagic flux via LC3-II accumulation in the presence/absence of lysosomal inhibitors (such as bafilomycin A1 or chloroquine) has been successfully applied *in vivo* in mammalian brain and other tissues, as well as in cell culture systems [51,52]. Systemic administration of these inhibitors to zebra finches, combined with western blot or immunofluorescence quantification of LC3-II in microdissected song nuclei could distinguish increased clearance from reduced debris generation. It may be helpful to develop methods to isolate and culture primary zebra finch astrocyte cultures, as *in vitro* approaches using lesion-mimetic stressors with and without CBD would allow more direct mechanistic dissection (e.g., flux assays with lysosomal blockade or p62/SQSTM1 turnover). Such combined *in vivo* and *in vitro* expansion would strengthen causal inference regarding CBD’s effects on astrocyte autophagic processes. The present lysosomal findings mechanistically extend our earlier observations of CBD-mediated reductions in SOD2, DHE fluorescence, and NRF2 pathway activation [20]. Activation of PPARG, a known transcriptional coordinator of antioxidant and metabolic programs in glia [53,54] offers a plausible convergent mechanism, as PPARG regulates both lysosomal function and mitochondrial turnover [55]. Future experiments using selective PPARG and/or TRPV antagonists may help to confirm pathway specificity.

CBD further enhanced astrocyte metabolic resilience by increasing expression of glutamine synthetase (GS) and glutamate-cysteine ligase modifier subunit (GCLM), the regulatory component of the rate-limiting enzyme in glutathione biosynthesis (Fig. 6). GS supports glutamate recycling and prevents excitotoxicity, while GCLM is a key cofactor for glutathione synthesis and downstream redox buffering: functions essential for maintaining synaptic homeostasis under injury stress [44,56,57]. These changes are consistent with the previously observed song circuit synaptic preservation and reduced excitotoxic vulnerability and suggest that CBD shifts astrocytes toward an oxidative stress–resistant, metabolically efficient phenotype [47,58] likely via enhanced GS/GCLM expression and NRF2-induced antioxidant responses. While NRF2 activation underpins antioxidant responses observed herein and in prior work, CBD likely engages complementary targets like PPARG for glial reactivity suppression and TLR4 inhibition for broader anti-inflammatory effects, as supported by recent mechanistic studies [59,60]. Future experiments with selective antagonists (e.g., GW9662 for PPARG) could help dissect these contributions in the zebra finch model. Although protein-level confirmation and assessment of the primary astrocytic glutamate transporter GLT1 also remains for future work, the transcriptional data (Figs 6 and 7) provide strong evidence that improved astrocytic management of glutamate/excitotoxic signaling supports the circuit stability required for sensorimotor relearning and vocal recovery.

Finally, CBD suppressed injury-induced astrocyte reactivity markers (C3, S100A10, aromatase; Fig. 7). C3 is typically associated with pro-inflammatory astrocytes, while S100A10 usually marks more homeostatic, anti-inflammatory astroglial states [27,61] and aromatase provides a general reactivity index [32,62]. Coordinated downregulation of all three transcripts, rather than selective promotion of a reparative phenotype (e.g., S100A10-expressing “A2-like” population), suggests that CBD prevents diversion of astrocytic resources into stress programs and sustains core homeostatic functions [47]. Such suppression is consistent with CBD-mediated inhibition of NFKB–driven transcription via PPARG and TRPV1 signaling [63,64] and aligns with the broad anti-inflammatory profile we documented previously. By limiting reactive astrogliosis, CBD appears to favor maintenance of the supportive astrocytic environment necessary for ongoing synaptic refinement and vocal circuit integrity.

Region-specific neuronal preservation in HVC (a site of adult neurogenesis) further suggests that CBD may enhance post-injury recruitment in addition to neuronal survival, potentially augmenting circuit repair [65,66]. Although proliferation markers (e.g., BrdU, DCX) were not assessed here, results raise the possibility that astrocyte-derived trophic support contributes to this effect [67,68].

Collectively, these findings establish astrocyte-targeted neuroprotection as a key mechanism by which CBD preserves song circuit integrity and accelerates recovery of a complex learned behavior after a focal TBI-like injury [16,20]. The zebra finch model, with discrete, well-mapped nuclei that support sensitive-period learning and lifelong sensorimotor maintenance, offers a powerful, translationally relevant platform for dissecting glial contributions to circuit resilience [69,70]. Many human sensorimotor skills (speech, language, manual dexterity) likewise depend on analogous specialized circuits vulnerable to traumatic disruption [65,71]. Our results therefore highlight the therapeutic potential of glial-directed interventions, with CBD or its downstream modulators as promising candidates for promoting functional recovery after CNS trauma [72,73].

## Conclusions

CBD preserves astrocyte viability, attenuates lysosomal stress/reactivity, and enhances metabolic/antioxidant support, stabilizing environments for synaptic homeostasis and sensorimotor relearning [42,47]. These provide a foundation for previously-established anti-inflammatory/synaptoprotective/behavioral effects [16,20]. The zebra finch vocal circuit advances understanding of neuroprotective mechanisms and glial-targeted therapy development [69,70,72].

## Supporting information

Supplemental Material for Cannabidiol Preserves Astrocyte Viability

## Acknowledgements

This work was supported by a grant from the U.S. Department of Defense (DoD) CDMRP #AZ220055 to K.S. and K.A.L. The authors also thank undergraduate laboratory members, Allie Kondracki, Mackenzie Lockard, Michelle Jacobs, Aishwarya Chinthakunta, and Ushaswini Mula, for their assistance with qRT-PCR experiments. We are grateful for Dr. Alessandro Didonna’s helpful review the manuscript.

## Abbreviations

AFP: anterior forebrain pathway
CBD: cannabidiol
CC3: cleaved caspase-3
DHE: dihydroethidium
GCLM: glutamate-cysteine ligase modifier subunit
GFAP: glial fibrillary acidic protein
GS: glutamine synthetase
HVC: (proper name; vocal premotor cortical-like nucleus)
IF: immunofluorescence
LAMP1: lysosomal-associated membrane protein 1
LC3: microtubule-associated protein 1A/1B-light chain 3
NeuN: neuronal nuclear protein
NRF2: nuclear factor erythroid 2-related factor 2
PBS: phosphate-buffered saline
PFA: paraformaldehyde
pNRF2: phosphorylated NRF2
PSD95: postsynaptic density protein 95
qRT-PCR: quantitative reverse transcription polymerase chain reaction
RA: robust nucleus of the arcopallium
REML: restricted maximum likelihood
SOD2: superoxide dismutase 2
TBI: traumatic brain injury
TMEM119: transmembrane protein 119
VGLUT2: vesicular glutamate transporter 2

