## Supplemental Material for Cannabidiol Preserves Astrocyte Viability for "Cannabidiol (CBD) Promotes Post-TBI Astrocyte Viability and Decreases Injury-Induced Glial Stress Responses Across Zebra Finch Song Control Nuclei"

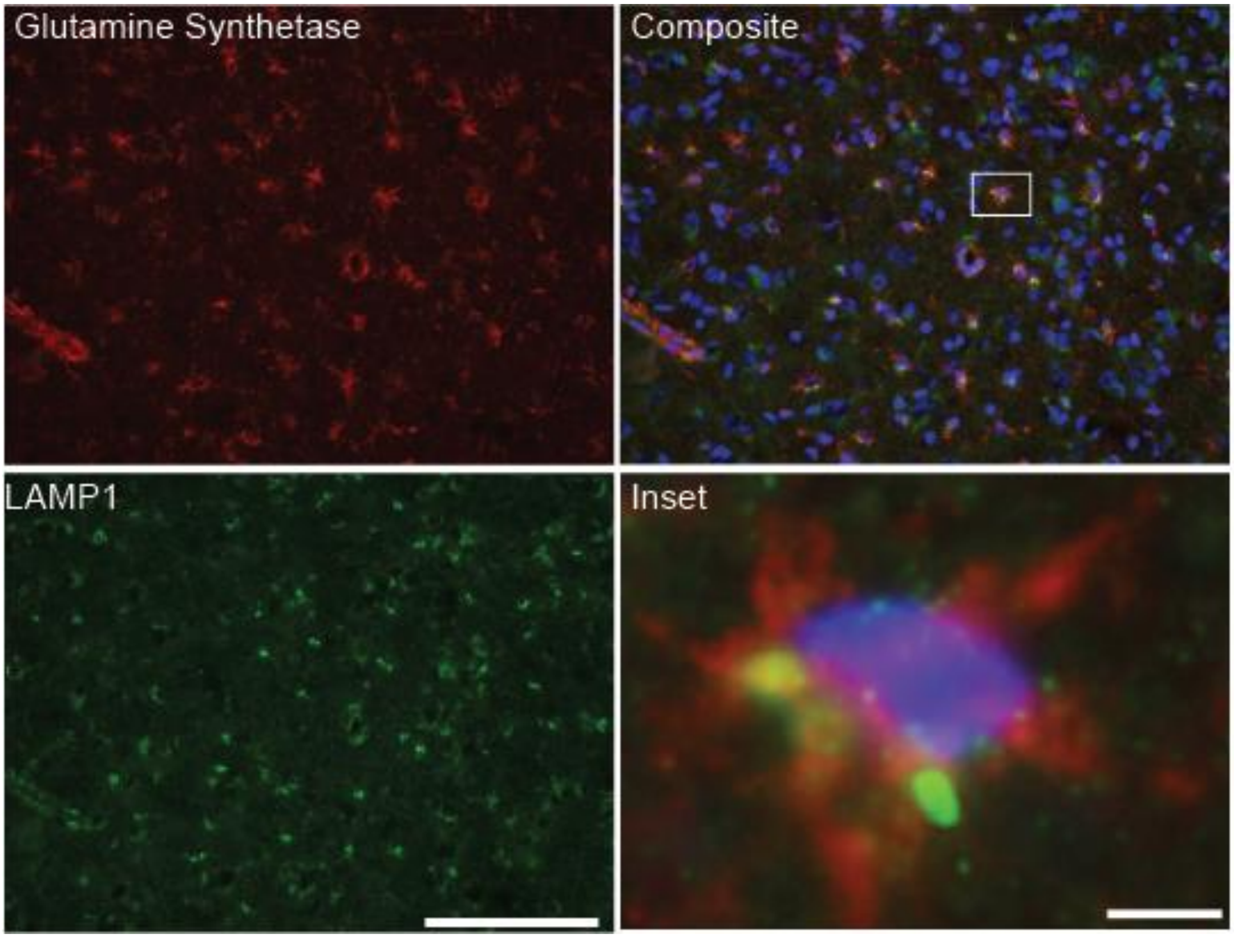


**Figure S1. Images of lysosomal LAMP1 localization in glutamine synthetase-positive astrocytes.** Representative images of glutamine synthetase (GS; red) LAMP1 (green) and Hoechst (blue) labeling in zebra finch brain tissue. GS immunoreactivity highlights astrocytic cell bodies and proximal processes (top left 40×) while LAMP1 labeling identifies lysosomal structures (bottom left 40×). The merged image (top right 40×) illustrates the spatial relationship between astrocytes and lysosomal compartments. The boxed region denotes the area shown at higher magnification (bottom right 263× digital zoom). The inset demonstrates the subcellular distribution of LAMP1 signal within GS-positive astrocytes with larger LAMP1-positive structures predominantly localized within GS-positive cells. Scale bars: 100 μm (top left top right bottom left) 10 μm (bottom right).

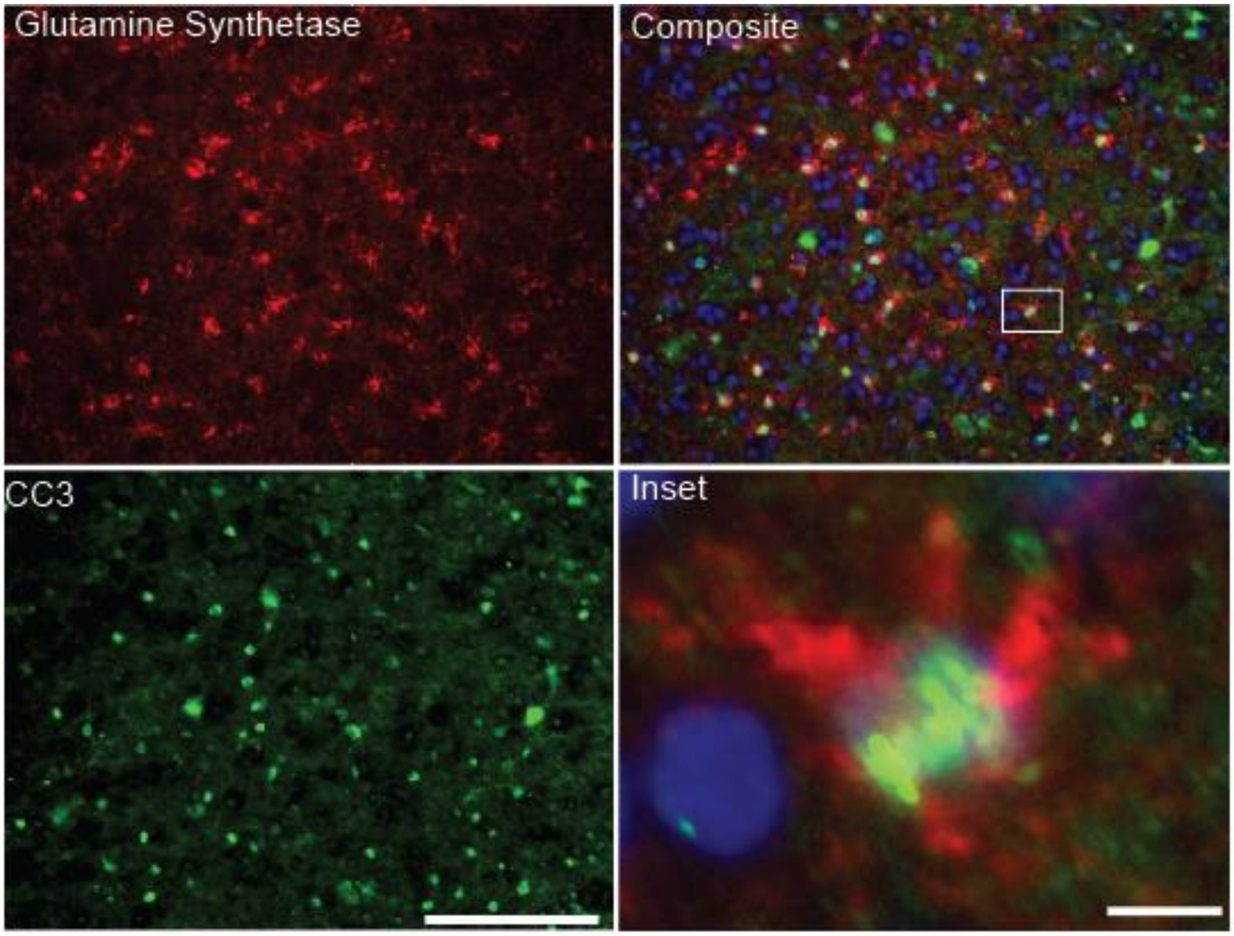


**Figure S2. Apoptotic (cleaved caspase-3-positive) glutamine synthetase-positive astrocytes.** Representative images of glutamine synthetase (GS; red) cleaved caspase-3 (CC3; green) and Hoechst (blue) labeling in zebra finch brain tissue. GS immunoreactivity highlights astrocytic cell bodies and proximal processes (top left 40×) while CC3 labeling identifies apoptotic cells (bottom left 40×). The merged image (top right 40×) illustrates the spatial relationship between CC3 expression and astrocytes. The boxed region denotes the area shown at higher magnification (bottom right 263× digital zoom). Most CC3-positive cells are localized within GS-positive astrocytes. Scale bars: 100 μm (top left top right bottom left) 10 μm (bottom right).

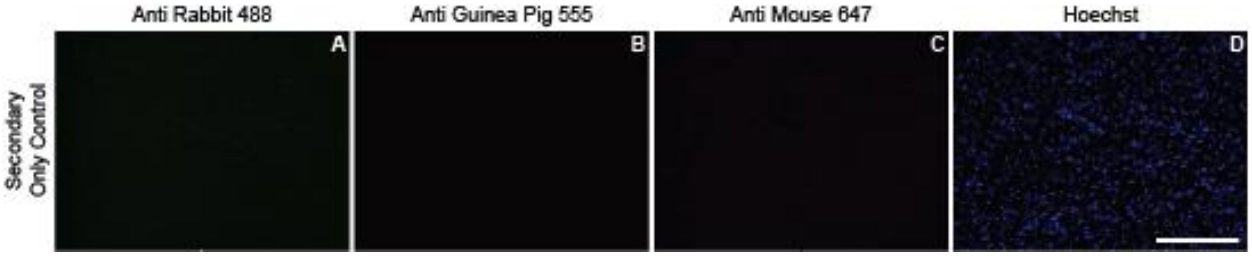


**Figure S3. Representative images of secondary-only control sections (20X, scale bar = 200 μm).** Representative images of secondary-only control sections (20× scale bar = 200 μm). Sections were incubated with secondary antibodies only including goat anti-rabbit Alexa Fluor 488 (Thermo Fisher Scientific Cat. #A-11008 1:500) (A) goat anti-guinea pig Alexa Fluor 555 (Thermo Fisher Scientific Cat. #A-21435 1:500) (B) and goat anti-mouse Alexa Fluor 647 (Thermo Fisher Scientific Cat. #A-21235 1:500) (C) with nuclei counterstained using Hoechst (D). Lipofuscin autofluorescence was quenched using TrueBlack Lipofuscin Autofluorescence Quencher (Cell Signaling Technology Cat. #92401 1:20) which reduces lipofuscin-associated and general background fluorescence. Minimal background fluorescence was observed across all channels confirming specificity of primary antibody labeling.

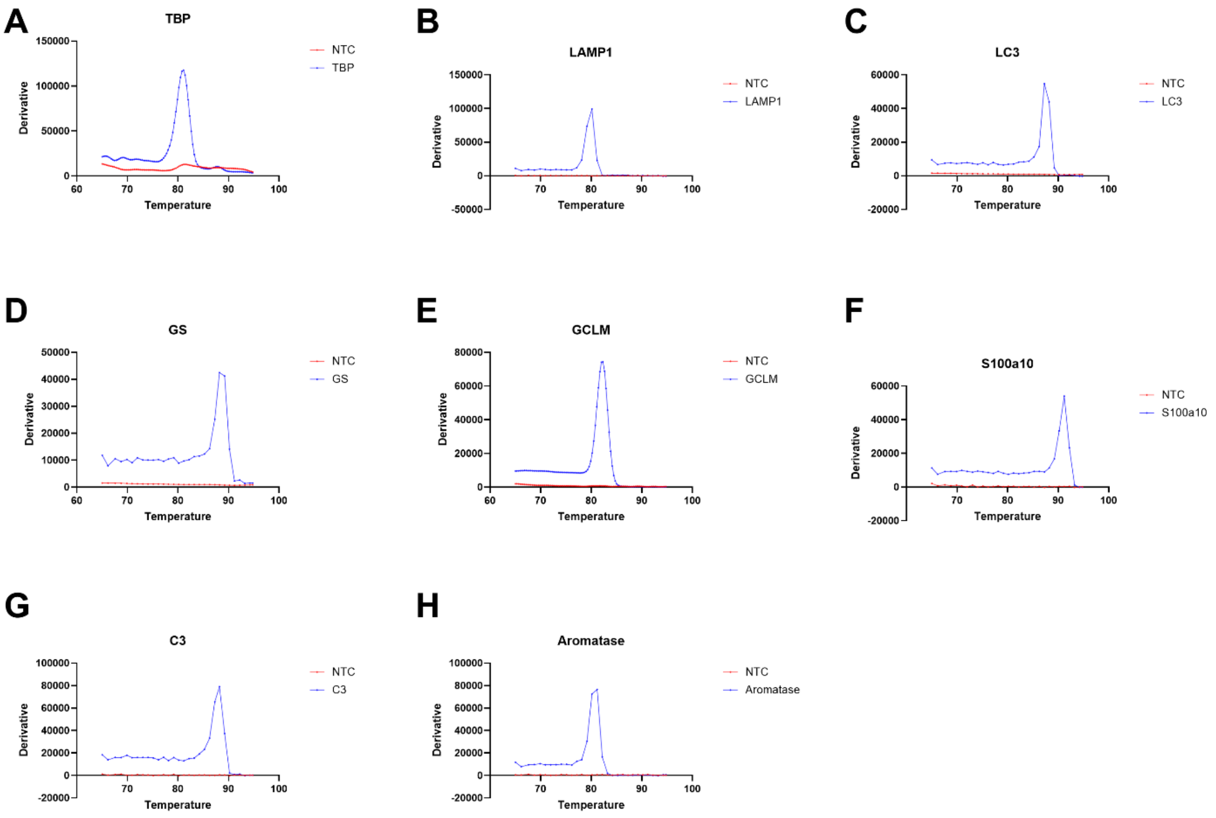


**Figure S4. Melt curve analysis confirms amplification specificity of qPCR targets.** Panels correspond to: (A) TATA binding protein (TBP), (B) LAMP1, (C) LC3, (D) GS, (E) GCLM, (F) S100a10, (G) C3, and (H) Aromatase. For each gene, a single, sharp melt peak was observed in reactions containing cDNA template, indicating amplification of a single specific product. In contrast, no distinct melt peak was observed in no-template controls (NTC), confirming the absence of non-specific amplification or primer-dimer formation

**Table S1. Summary of main effects and interactions from mixed-effects models ANOVAs.**

| Figure | Effect | F (DFn, DFd) | p-value |
|---|---|---|---|
| 4D | Region | F(2, 12) = 5.333 | 0.022 |
| 4D | Lesion condition | F(1, 6) = 11.36 | 0.015 |
| 4D | Treatment | F(1, 6) = 42.47 | 0.0006 |
| 4D | Region × Lesion condition | F(2, 12) = 0.3589 | 0.7057 |
| 4D | Region × Treatment | F(2, 12) = 0.8693 | 0.4441 |
| 4D | Lesion condition × Treatment | F(1, 6) = 8.026 | 0.0298 |
| 4D | Region × Lesion condition × Treatment | F(2, 12) = 5.793 | 0.0173 |
| 3D | Region | F(2, 12) = 30.69 | <0.0001 |
| 3D | Lesion condition | F(1, 6) = 0.4152 | 0.5432 |
| 3D | Treatment | F(1, 6) = 48.33 | 0.0004 |
| 3D | Region × Lesion condition | F(2, 12) = 9.122 | 0.0039 |
| 3D | Region × Treatment | F(2, 12) = 8.061 | 0.006 |
| 3D | Lesion condition × Treatment | F(1, 6) = 1.994 | 0.2076 |
| 3D | Region × Lesion condition × Treatment | F(2, 12) = 2.347 | 0.1379 |
| 3F | Region | F(2, 12) = 252.8 | <0.0001 |
| 3F | Lesion condition | F(1, 6) = 15.65 | 0.0075 |
| 3F | Treatment | F(1, 6) = 2.078 | 0.1995 |
| 3F | Region × Lesion condition | F(2, 12) = 6.610 | 0.0116 |
| 3F | Region × Treatment | F(2, 12) = 10.61 | 0.0022 |
| 3F | Lesion condition × Treatment | F(1, 6) = 0.1862 | 0.6811 |
| 3F | Region × Lesion condition × Treatment | F(2, 12) = 3.044 | 0.0853 |
| 3G | Region | F(2, 12) = 5.755 | 0.0177 |
| 3G | Lesion condition | F(1, 6) = 8.513 | 0.0267 |
| 3G | Treatment | F(1, 6) = 1.853 | 0.2223 |
| 3G | Region × Lesion condition | F(2, 12) = 9.148 | 0.0039 |
| 3G | Region × Treatment | F(2, 12) = 4.351 | 0.0379 |
| 3G | Lesion condition × Treatment | F(1, 6) = 3.154 | 0.1261 |
| 3G | Region × Lesion condition × Treatment | F(2, 12) = 5.408 | 0.0212 |
| 3E | Region | F(2, 12) = 15.22 | 0.0005 |
| 3E | Lesion condition | F(1, 6) = 120.7 | <0.0001 |
| 3E | Treatment | F(1, 6) = 39.35 | 0.0008 |
| 3E | Region × Lesion condition | F(2, 12) = 14.03 | 0.0007 |
| 3E | Region × Treatment | F(2, 12) = 0.1776 | 0.8395 |
| 3E | Lesion condition × Treatment | F(1, 6) = 24.17 | 0.0027 |
| 3E | Region × Lesion condition × Treatment | F(2, 12) = 1.176 | 0.3417 |
| 6A | Region | F(2, 18) = 11.23 | 0.0007 |
| 6A | Treatment | F(1, 18) = 51.09 | <0.0001 |
| 6A | Region × Treatment | F(2, 18) = 19.29 | <0.0001 |
| 6B | Region | F(2, 18) = 15.92 | 0.0001 |
| 6B | Treatment | F(1, 18) = 34.23 | <0.0001 |
| 6B | Region × Treatment | F(2, 18) = 16.51 | <0.0001 |
| 5A | Region | F(2, 18) = 162.9 | <0.0001 |
| 5A | Treatment | F(1, 18) = 30.12 | <0.0001 |
| 5A | Region × Treatment | F(2, 18) = 19.13 | <0.0001 |
| 5B | Region | F(2, 18) = 117.9 | <0.0001 |
| 5B | Treatment | F(1, 18) = 10.40 | 0.0047 |
| 5B | Region × Treatment | F(2, 18) = 27.50 | <0.0001 |
| 7C | Region | F(2, 12) = 70.95 | <0.0001 |
| 7C | Treatment | F(1, 6) = 24.82 | 0.0025 |
| 7C | Region × Treatment | F(2, 12) = 47.72 | <0.0001 |
| 7A | Region | F(2, 12) = 37.39 | <0.0001 |
| 7A | Treatment | F(1, 6) = 5.312 | 0.0607 |
| 7A | Region × Treatment | F(2, 12) = 9.643 | 0.0032 |
| 7B | Region | F(2, 12) = 84.37 | <0.0001 |
| 7B | Treatment | F(1, 6) = 14.71 | 0.0086 |
| 7B | Region × Treatment | F(2, 12) = 0.8347 | 0.4577 |

### Table S2. Region-specific post hoc comparisons across treatment and lesion conditions

| Figure | Region | Comparison | Mean Difference | 95% CI | Adjusted p-value |
|---|---|---|---|---|---|
| 3D | HVC | VEH Un-lesioned vs. VEH Lesioned | -1.217 | -3.735 to 1.301 | 0.4237 |
| 3D | HVC | CBD Un-lesioned vs. CBD Lesioned | -4.131 | -6.649 to -1.612 | 0.0025 |
| 3D | HVC | VEH Un-lesioned vs. CBD Un-lesioned | -2.414 | -4.932 to 0.1047 | 0.0606 |
| 3D | HVC | VEH Lesioned vs. CBD Lesioned | -5.327 | -7.845 to -2.809 | 0.0003 |
| 3D | RA | VEH Un-lesioned vs. VEH Lesioned | -0.3385 | -5.167 to 4.490 | 0.9752 |
| 3D | RA | CBD Un-lesioned vs. CBD Lesioned | 1.487 | -3.341 to 6.315 | 0.6366 |
| 3D | RA | VEH Un-lesioned vs. CBD Un-lesioned | -9.619 | -14.60 to -4.635 | 0.0007 |
| 3D | RA | VEH Lesioned vs. CBD Lesioned | -7.793 | -12.78 to -2.809 | 0.0036 |
| 3D | Area X | VEH Un-lesioned vs. VEH Lesioned | 5.016 | 1.275 to 8.757 | 0.0147 |
| 3D | Area X | CBD Un-lesioned vs. CBD Lesioned | 1.351 | -2.390 to 5.092 | 0.5462 |
| 3D | Area X | VEH Un-lesioned vs. CBD Un-lesioned | -0.6725 | -4.503 to 3.158 | 0.8858 |
| 3D | Area X | VEH Lesioned vs. CBD Lesioned | -4.338 | -8.169 to -0.5068 | 0.0269 |
| 3E | HVC | VEH Un-lesioned vs. VEH Lesioned | -32.81 | -43.26 to -22.37 | 0.0002 |
| 3E | HVC | CBD Un-lesioned vs. CBD Lesioned | -9.774 | -20.22 to 0.6739 | 0.0640 |
| 3E | HVC | VEH Un-lesioned vs. CBD Un-lesioned | 6.407 | -4.772 to 17.58 | 0.3096 |
| 3E | HVC | VEH Lesioned vs. CBD Lesioned | 29.45 | 18.27 to 40.63 | <0.0001 |
| 3E | RA | VEH Un-lesioned vs. VEH Lesioned | -21.25 | -42.57 to 0.0655 | 0.0498 |
| 3E | RA | CBD Un-lesioned vs. CBD Lesioned | -6.914 | -28.23 to 14.41 | 0.6085 |
| 3E | RA | VEH Un-lesioned vs. CBD Un-lesioned | 7.94 | -10.54 to 26.42 | 0.5018 |
| 3E | RA | VEH Lesioned vs. CBD Lesioned | 22.28 | 3.802 to 40.76 | 0.019 |
| 3E | Area X | VEH Un-lesioned vs. VEH Lesioned | -53.11 | -63.75 to -42.47 | <0.0001 |
| 3E | Area X | CBD Un-lesioned vs. CBD Lesioned | -24.23 | -34.87 to -13.59 | 0.0010 |
| 3E | Area X | VEH Un-lesioned vs. CBD Un-lesioned | 0.515 | -10.05 to 11.08 | 0.9906 |
| 3E | Area X | VEH Lesioned vs. CBD Lesioned | 29.4 | 18.83 to 39.96 | <0.0001 |
| 3F | HVC | VEH Un-lesioned vs. VEH Lesioned | -12.69 | -26.24 to 0.8679 | 0.0638 |
| 3F | HVC | CBD Un-lesioned vs. CBD Lesioned | -13.45 | -27.01 to 0.1033 | 0.0515 |
| 3F | HVC | VEH Un-lesioned vs. CBD Un-lesioned | -14.02 | -27.98 to -0.0695 | 0.0489 |
| 3F | HVC | VEH Lesioned vs. CBD Lesioned | -14.79 | -28.74 to -0.8342 | 0.0378 |
| 3F | RA | VEH Un-lesioned vs. VEH Lesioned | 6.07 | -4.046 to 16.19 | 0.2364 |
| 3F | RA | CBD Un-lesioned vs. CBD Lesioned | -5.677 | -15.79 to 4.439 | 0.2739 |
| 3F | RA | VEH Un-lesioned vs. CBD Un-lesioned | 4.293 | -6.305 to 14.89 | 0.5396 |
| 3F | RA | VEH Lesioned vs. CBD Lesioned | -7.455 | -18.05 to 3.143 | 0.1859 |
| 3F | Area X | VEH Un-lesioned vs. VEH Lesioned | -14.36 | -28.13 to -0.6018 | 0.0624 |
| 3F | Area X | CBD Un-lesioned vs. CBD Lesioned | -6.989 | -20.75 to 6.773 | 0.3335 |
| 3F | Area X | VEH Un-lesioned vs. CBD Un-lesioned | -0.8486 | -13.03 to 11.34 | 0.9809 |
| 3F | Area X | VEH Lesioned vs. CBD Lesioned | 6.526 | -5.658 to 18.71 | 0.3544 |
| 3G | HVC | VEH Un-lesioned vs. VEH Lesioned | -14.51 | -25.04 to -3.984 | 0.0130 |
| 3G | HVC | CBD Un-lesioned vs. CBD Lesioned | -2.587 | -13.12 to 7.942 | 0.7446 |
| 3G | HVC | VEH Un-lesioned vs. CBD Un-lesioned | 0.1046 | -9.089 to 9.298 | 0.9995 |
| 3G | HVC | VEH Lesioned vs. CBD Lesioned | 12.03 | 2.836 to 21.22 | 0.0117 |
| 3G | RA | VEH Un-lesioned vs. VEH Lesioned | -1.634 | -8.937 to 5.670 | 0.7816 |
| 3G | RA | CBD Un-lesioned vs. CBD Lesioned | -1.063 | -8.366 to 6.241 | 0.8988 |
| 3G | RA | VEH Un-lesioned vs. CBD Un-lesioned | -0.1096 | -6.634 to 6.415 | 0.9989 |
| 3G | RA | VEH Lesioned vs. CBD Lesioned | 0.4611 | -6.063 to 6.985 | 0.9803 |
| 3G | Area X | VEH Un-lesioned vs. VEH Lesioned | -2.082 | -4.921 to 0.7570 | 0.1408 |
| 3G | Area X | CBD Un-lesioned vs. CBD Lesioned | -0.7842 | -3.623 to 2.055 | 0.6919 |
| 3G | Area X | VEH Un-lesioned vs. CBD Un-lesioned | -1.063 | -3.541 to 1.415 | 0.5030 |
| 3G | Area X | VEH Lesioned vs. CBD Lesioned | 0.2352 | -2.243 to 2.713 | 0.9649 |

| Figure | Region | Comparison | Mean Difference | 95% CI | Adjusted p-value |
|---|---|---|---|---|---|
| 4D | HVC | VEH Un-lesioned vs. VEH Lesioned | -0.3163 | -2.096 to 1.463 | 0.8540 |
| 4D | HVC | CBD Un-lesioned vs. CBD Lesioned | -1.815 | -3.595 to -0.0359 | 0.0463 |
| 4D | HVC | VEH Un-lesioned vs. CBD Un-lesioned | 4.86 | 2.407 to 7.313 | 0.0006 |
| 4D | HVC | VEH Lesioned vs. CBD Lesioned | 3.361 | 0.9073 to 5.814 | 0.0088 |
| 4D | RA | VEH Un-lesioned vs. VEH Lesioned | 1.347 | -1.050 to 3.745 | 0.2729 |
| 4D | RA | CBD Un-lesioned vs. CBD Lesioned | -2.677 | -5.074 to -0.2797 | 0.0324 |
| 4D | RA | VEH Un-lesioned vs. CBD Un-lesioned | 4.973 | 2.649 to 7.297 | 0.0003 |
| 4D | RA | VEH Lesioned vs. CBD Lesioned | 0.9488 | -1.375 to 3.273 | 0.5347 |
| 4D | Area X | VEH Un-lesioned vs. VEH Lesioned | -1.501 | -3.383 to 0.3807 | 0.1094 |
| 4D | Area X | CBD Un-lesioned vs. CBD Lesioned | -0.9359 | -2.818 to 0.9458 | 0.3464 |
| 4D | Area X | VEH Un-lesioned vs. CBD Un-lesioned | 3.051 | 1.073 to 5.028 | 0.0039 |
| 4D | Area X | VEH Lesioned vs. CBD Lesioned | 3.616 | 1.638 to 5.594 | 0.0011 |
| 5A | HVC | VEH Lesioned vs. CBD Lesioned | 0.7235 | 0.4153 to 1.0320 | <0.0001 |
| 5A | RA | VEH Lesioned vs. CBD Lesioned | 0.6066 | 0.2984 to 0.9148 | 0.0002 |
| 5A | Area X | VEH Lesioned vs. CBD Lesioned | -0.2166 | -0.5248 to 0.0916 | 0.2238 |
| 5B | HVC | VEH Lesioned vs. CBD Lesioned | 0.6258 | 0.3482 to 0.9034 | <0.0001 |
| 5B | RA | VEH Lesioned vs. CBD Lesioned | 0.3913 | 0.1137 to 0.6689 | 0.0048 |
| 5B | Area X | VEH Lesioned vs. CBD Lesioned | -0.4278 | -0.7054 to -0.1503 | 0.0022 |
| 6A | HVC | VEH Lesioned vs. CBD Lesioned | -1.035 | -1.505 to -0.5655 | <0.0001 |
| 6A | RA | VEH Lesioned vs. CBD Lesioned | -1.329 | -1.798 to -0.8586 | <0.0001 |
| 6A | Area X | VEH Lesioned vs. CBD Lesioned | 0.1529 | -0.3170 to 0.6228 | 0.7874 |
| 6B | HVC | VEH Lesioned vs. CBD Lesioned | -0.9411 | -1.268 to -0.6139 | <0.0001 |
| 6B | RA | VEH Lesioned vs. CBD Lesioned | -0.3868 | -0.7140 to -0.0596 | 0.0180 |
| 6B | Area X | VEH Lesioned vs. CBD Lesioned | 0.0677 | -0.2595 to 0.3949 | 0.9324 |
| 7A | HVC | VEH Lesioned vs. CBD Lesioned | 1.219 | 0.5593 to 1.8790 | 0.0004 |
| 7A | RA | VEH Lesioned vs. CBD Lesioned | 0.0095 | -0.6505 to 0.6694 | >0.9999 |
| 7A | Area X | VEH Lesioned vs. CBD Lesioned | -0.1713 | -0.8312 to 0.4886 | 0.8774 |
| 7B | HVC | VEH Lesioned vs. CBD Lesioned | 1.549 | 0.2904 to 2.8070 | 0.0136 |
| 7B | RA | VEH Lesioned vs. CBD Lesioned | 1.457 | 0.1988 to 2.7160 | 0.0207 |
| 7B | Area X | VEH Lesioned vs. CBD Lesioned | 0.8456 | -0.4129 to 2.1040 | 0.2564 |
| 7C | HVC | VEH Lesioned vs. CBD Lesioned | 2.268 | 1.698 to 2.838 | <0.0001 |
| 7C | RA | VEH Lesioned vs. CBD Lesioned | -0.2136 | -0.7835 to 0.3563 | 0.7087 |
| 7C | Area X | VEH Lesioned vs. CBD Lesioned | 0.1575 | -0.4124 to 0.7274 | 0.8565 |

**Table S3. Primers used for quantitative RT-PCR**

| Gene Symbol | Accession Number | Forward Primer | Reverse Primer |
|---|---|---|---|
| TBP | XM_030268321.4 | TCACACACCAGCAGTTCAGC | CTGCTCGTACTTTAGCACCAGTTA |
| GS | XM_030279644.4 | AATCTGCTCCCCAGCTTCCC | TCAGCCTTTCCAGAGCAGATCA |
| GCLM | XM_030279191.4 | GACTGAGTGGAGCTCAAAGGT | CCAGGTCAACTGCGTCTCTG |
| LAMP1 | XM_002191571.5 | TGCATCAACTCCAAGCACATC | AGAGACCCTATCTTCAGCACATTC |
| LC3B | NM_001245293.2 | AGCTTCCAGTTTTGGATAAGACC | TTCTCGCTCTCGTACACCTC |
| CYP19A1 | AH008871.2 | CTCCACCGACAAAAATCCAC | GGGTTTCCAGGACCATCTTT |
| S100a10 | XM_030291396.4 | TCATGGACAAGGAGTTCCCG | TGCTGCACGAAGTAGTCGTT |
| C3 | XM_072920066.1 | GGCTCTCCCAGCAGAACTAC | AGAACCGACACGTCCAACTC |
